# Spatial dissection of cellular heterogeneity and functional crosstalk in the tumor microenvironments of gastric cancers

**DOI:** 10.1101/2023.12.24.573275

**Authors:** Sung Hak Lee, Dagyeong Lee, Junyong Choi, Hye Jeong Oh, In-Hye Ham, Daeun Ryu, Seo-Yeong Lee, Dong-Jin Han, Sunmin Kim, Seungho Lee, Hoon Hur, Tae-Min Kim

## Abstract

Spatial transcriptomics offers unprecedented challenges in characterizing tumor ecosystems, encompassing tumor cells and their surrounding tumor microenvironments (TME) along with cellular interactions therein. Here, we conducted Visium-based spatial transcriptomics on nine primary gastric cancers (GC) to translate the spatial organization of GC ecosystems into the functional landscape of cellular crosstalk across malignant, stromal and immune cells. We identified three distinct GC subtypes of 3 immunogenic, 3 epithelial and 3 stromal GCs with elevated infiltration of immune, malignant cells and fibroblasts reflecting the heterogeneity of the cell types and their interactions in GC TME. GC spatial architecture was further delineated into 6 regional compartments with varying degree of TME infiltration, among which the fibroblast-enriched TME was associated with the upregulation of epithelial-to-mesenchymal transformation (EMT) and immune response in malignant and immune cells, respectively. Cell type-specific transcriptional dynamics representing cellular crosstalk between TME cells were further identified, e.g., the infiltration of malignant and endothelial cells promote the cellular proliferations of TME cells whereas the fibroblasts and immune cells are associated with pro-cancer and anti-cancer immunity, respectively. Ligand-receptor analysis further revealed that *CCL2*-expressing fibroblasts promote the pro-cancer signaling of immune cells including JAK-STAT3 signaling and inflammatory response in tumor-infiltration macrophages. The *CCL2+*fibroblasts and macrophages with activated JAK-STAT3 signals are co-localized in spatial GC architecture and their co-abundance was associated with unfavorable clinical outcomes. Taken together, GC spatial transcriptomes can be regionally and functionally delineated into functional cellular crosstalk involving multiple cell types with potential clinical relevance, particularly the interaction between *CCL2+* fibroblasts and JAK-STAT3+macrophages contributing to cancer-favoring immune contexture in GC TME.

## Introduction

Gastric cancers (GC) are major global concerns with their high cancer-related morbidity and mortality (1). Recently, large-scaled sequencing efforts have revealed universal GC driver genes such as *TP53* mutations (2,3) and GC-specific mutations such as *ROHA* mutations (4,5) that have improved our understanding into the mechanisms underlying GC development. While the mutational configuration may guide the selection of molecular targets, the effect is only modest (6) and the currently approved therapeutic regimen is *Her2* positivity and its targeting Trastuzumab (7). The standard therapeutic modalities of GCs – the early intervention and surgical resection along with cytotoxic agents-based chemotherapy such as 5-FU (fluorouracil) – have been not much improved for decades. Although the emergence of immune checkpoint inhibitors may revolutionize the GC therapeutic modalities; the selection of patients with benefits to the therapy is largely hampered with the well-known tumor heterogeneity (8) and lack of appropriate markers (9)

It is appreciated that the disease heterogeneity responsible for the disease initiation, maintenance and also the progression are largely attributed to the transcriptional and cellular heterogeneity. To cope with the heterogeneity, bulk-level RNA sequencing (RNA-seq) has been utilized for GC subclassification (‘molecular taxonomy’) leading to the 4 GC subtypes by the ACRG report (10), but the data does not correctly deal with the tumor heterogeneity (11). Recently developed single cell RNA sequencing (scRNA-seq) technologies directly evaluate the transcriptional heterogeneity of tissues at single cell resolution also facilitating the delineation of subtypes of major cell types in tumor microenvironments (TME). For GC, Zhang et al. analyzed the scRNA-seq of premalignant GC (12). Sathe (13) and Zhang (14) analyzed the scRNA-seq of GC identifying cellular compositions, gene-level markers and evidence of cellular interactions in terms of ligand-receptor pairs underlying the GC biology. Currently available, large scRNA-seq data (15) may also serve as valuable resources in cellular heterogeneity of GC facilitating the identification of cellular subpopultations associated with particular phenotypes of interests such as the drug resistance, metastatic risk and cellular plasticity.

Nevertheless, bulk-level RNA-seq and scRNA-seq techniques lack the ability to preserve the anatomical and spatial information of tissues or cells under investigation since the specimens are dissociated during the experimental procedures. In a previous study (16), we performed tumor depth-aware scRNA-seq leveraging the spatial information by separately analyzing layer-specific scRNA-seq of diffuse-type GCs (i.e., normal tissues and superficial-*vs*.-deep tumor layers). By assuming that the transitions from superficial to deep layers reflect the tumor progression, we were able to identify cellular subpopulations exhibiting differential expression along the transitions. Recently, high-resolution mapping of the spatial organization of cells become possible generating thousands of transcriptome profiles conserved for their anatomical contexts of tissue architecture per sample (17). Solid tumors, including GC, are ideal candidates for spatial transcriptomics since the development and progression of tumors occur in the complex cellular milieu of malignant, immune and stromal cells in TME (18). In addition to spatially unreveling the cellular heterogeneity, these technologies have the potential to interrogate the spatial relationship between malignant cells and TME cells and to explore the functional consequences in terms of spatial patterns of oncogenic crosstalks.

In this study, we employed barcode-based Visium spatial transcriptome technology (19) for nine GC specimens (GC1 to GC9). We first established three GC subtypes based on cell abundance to ensure that the histology of GC is robustly represented by cellular composition of given GC tissue. The deconvoluted spot-level cell abundance was then employed to further categorize six distinct regions with differential level of TME cell infiltrations in GC tissue architecture. Next, we applied techniques to infer gene expression specific to cell types within these regions to identify the transcriptional behavior of malignant cells, fibroblasts, endothelial and immune cells in relation to TME infiltrations. Furthermore, we established landscape of functional cellular crosstalk representing functional consequences of cellular interactions between regulator and target cells. Among the interactions, we highlighted that *CCL2+* fibroblasts regulate the JAK-STAT signaling of macrophages as key immunomodulatory impact of the fibroblast infiltration in GC TME.

## Results

### Spatial cellular maps of three GC subtypes

Nine GC surgical specimens (all surgically removed primary GCs) were obtained from 9 patients and were subject to spatial transcriptomics by Visium technologies. Clinicoopathological features of nine GC patients are available in **Supplementary Table 1/patient**. Overall, spatial transcriptomics data were obtained yielding data from 1882 - 4274 spots per histologic section of 9 cases. The median number of spots were 3,491 spots with a median depth of unique molecular identifier (UMI) of 3,491 genes per spot (**Supplementary Table 2/sequencing**). The spot-level cell deconvolution was performed with respect to 11 cell types as available in our previous scRNA-seq data of five GC cases (16). Epithelial cells were further divided into those on tumor and normal regions (malignant and normal epithelium, respectively) so that the deconvoluted cellular abundance per spot was examined for 12 cell types in total. The summarization of deconvoluted cellular abundance, categorized on a case-by-case basis, is shown for nine GC spatial transcriptomics in **Fig. 1A**. Notably, while the cellular compositions reveal evident heterogeneity, the clustering of the deconvoluted cellular abundance distinguishes three GC subtypes: ‘immunogenic’, ‘epithelial’, and ‘fibrotic’. These subtypes are largely consistent with the Lauren’s classification, e.g., intestinal type GCs were either classified into immunogenic or epithelial GC subtypes while all the fibrotic GCs to the diffuse type. The main cellular components of three epithelial GCs (GC3, GC5 and GC7) are malignant or normal epithelial cells. Three immunogenic GCs were either microsatellite unstable (microsatellite instability or MSI-positive; GC1) or Epstein-Barr virus-positive GCs (GC2 and GC8), consistent with their high-level of tumor infiltrating immune cells as previously reported (20). CD4+ and CD8+ T cells are major immune cell components in these GC types. Among 5 diffuse types of GCs, three (GC4, GC6 and GC9) were annotated as fibrotic GCs with high-level of tumor infiltrating fibroblasts.

**Figure 1.**
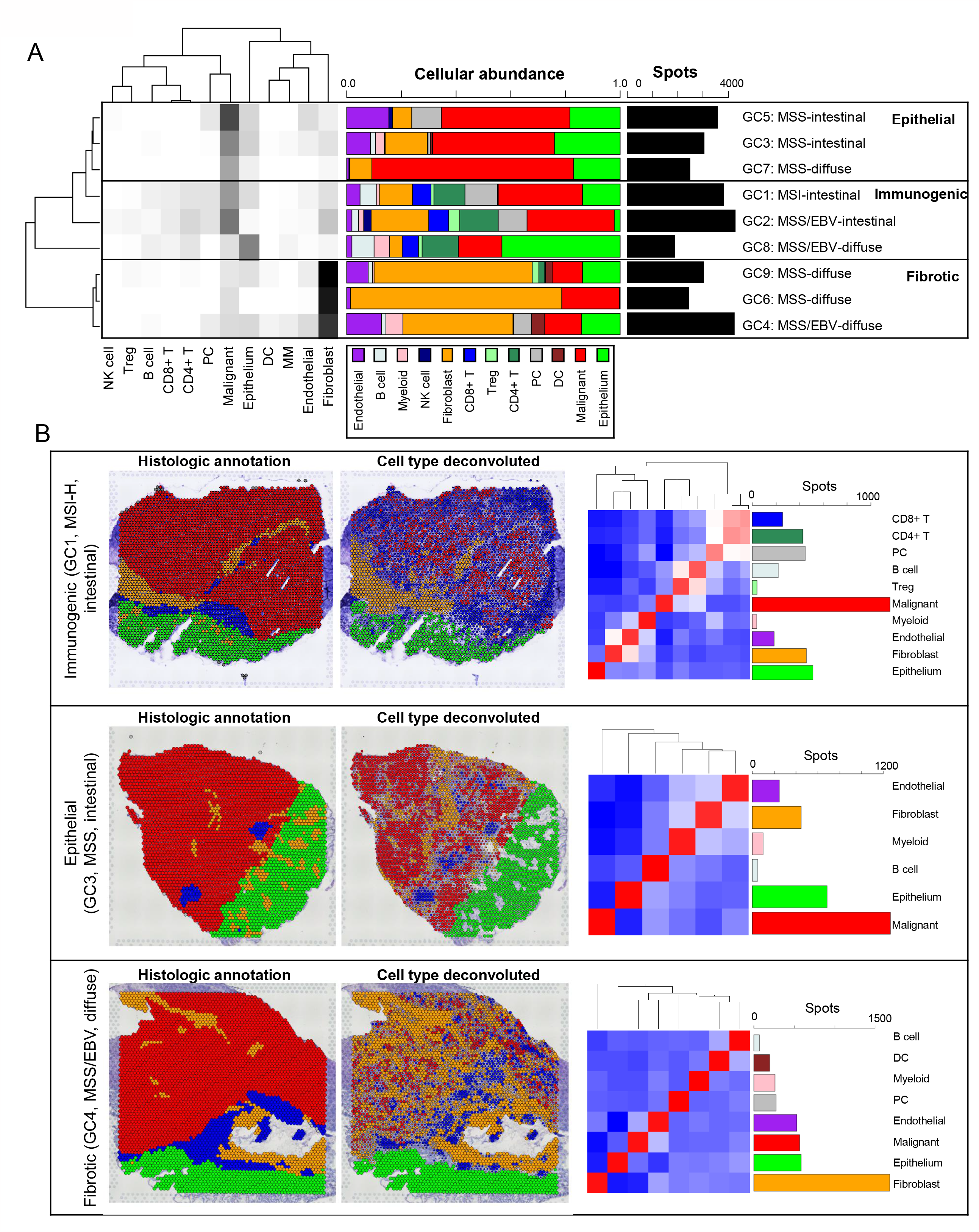
Classification of three GC subtypes and their spatial cellular architecture. (A) A heatmap (on the left) illustrating the cellular abundance of 12 cell types for nine GC cases (GC1 to GC9) along with barplots (in the middle) depicting the cellular abundance for each case. Three distinct subtypes: epithelial, immunogenic, and fibrotic GC subtypes were distinguished with the prevalence of malignant cells, immune cells and fibroblasts, respectively. (B) For three representative cases - immunogenic (GC1), epithelial (GC3), and fibrotic (GC4) GCs - histological annotations as determined by pathologists (left) are shown with the dominant cell types of individual spots, as identified through deconvolution (middle). A correlative heatmap also demonstrates the spatial-level correlation of cellular abundances (right).

Spatial cellular maps for selected examples across three GC categories (GC1/GC3/GC4 representing immunogenic, epithelial and fibrotic GCs, respectively) are depicted in **Figure 1B**. Two types of spatial maps are represented: the histology-based regional annotation and deconvoluted spot-level cellular abundances highlighting cell types with predominant presence or the highest abundances per individual spot. Additionally, the abundance of individual cell types are presented along correlative heatmaps depicting the level of correlation across spots. The spatial maps of the remaining cases are shown in **Supplementary Fig. 1**. In the instance of GC1 (immunogenic, MSI-H, intestinal type), a distinct regional differentiation between normal epithelial and stromal cells, was observed. While malignant cells exhibited substantial immune cell infiltration, the tumor stroma was primarily composed of fibroblasts and endothelial cells, with their concurrent presence suggestive of fibrovascular niche (21). The major immune cell components of GC1, including T cells (CD4+ and CD8+) and plasma cells (PCs), were regionally coincident as shown in the correlative heatmap. Furthermore, a regional association of B cells and Tregs with malignant cells was observed, while myeloid cells (monocyte-macrophages, MM) were associated with tumor stroma. Therefore, immunogenic GCs exhibit an association of fibroblasts and endothelial cells, forming a fibrovascular niche also connected to myeloid lineage cells. Immune cells can be classified into two categories based on their regional concordance with malignant cells: those infiltrating the tumor (Tregs and B cells) and those that are exclusive (T cells and PCs). The other immunogenic GCs (GC2 and GC8, **Supplementary Fig. 1**) demonstrated similar spatial characteristic of GC1. Epithelial GC (GC3, MSS, intestinal type) also displays evidence of a fibrovascular niche, as indicated by the regional conjuction of fibroblasts and endothelial cells in the tumor regions. Despite a relative depletion of T cells and PCs in epithelial GCs, MM cells and B cells were still present, establishing immune cell-infiltrating niches in tumor regions. In a fibrotic GC (GC4, MSS, diffuse type), the tumor regions displayed a prominent and vast infiltration of fibroblasts, demonstrating a distinct pattern in contrast to immunogenic or epithelial GCs. A considerable presence of tumor infiltrating immune cells was noted, primarily composed of PCs, B cells, MM cells and DCs, albeit without T cells. The remaining epithelial (GC5 and GC7) and fibrotic GCs (GC6 and GC9) are shown in **Supplementary Fig. 1**. Consequently, the cellular abundance in GC microenvironments can be classified into at least three GC subtypes. The spatially resolved cellular abundance offers insights into the major cell types present and their spatial relationships in GC TME architectures.

### Spatially resolved cellular and transcriptional dynamics in GC tissue architecture

To regionally dissect the spatial cellular heterogeneity in GC TME, we employed a method for the regional categorization of spots taking into accounts the varying cellular constituents and their regional adjacency. For individual spots, 6 spots directly adjacent to the given spots (referred to as ‘N1’ spots) and the 12 spots adjacent to the N1 spots (referred to as ‘N2’ spots) were examined for the cellular abundance of 12 cell types, The cellular abundance of the original spots of interests (‘S’ spots) were jointly analyzed with their matching N1 and N2 spots to gain insights into the local cellular architecture of the TME, with the details illustrated in **Supplementary Fig. 2**. The hierarchical clustering of all the spots of nine GCs (*n* = 29,808) with respect to the cellular abundance of S, N1 and N2 spots revealed six regional categories, each with varying cellular components and the extent of TME infiltration (**Fig. 2A**). The resulting categories, termed “Cellular Composite Zones” (CCZs) were annotated based on their cellular composition and the dominant cell types. For instance, zones primarily consisting of normal and malignant epithelial cells were designated as ‘Epithelial’ and ‘Malignant-dominant’ CCZs, respectively. Those with a prevalence of fibroblasts were labeled ‘Fibroblast-dominant’. The remaining categories, marked by the infiltration of diverse cell types, were annotated as ‘Malignant-infiltrated’, ‘Fibroblast-infiltrated’, and ‘Immune-dominant’, each indicative of the respective cellular dynamics within the TME. **Figure 2B** delineates the cellular composition within the six identified CCZs and correlates these zones to three distinct GC subtypes. A detailed spatial CCZ map for a representative immunogenic GC case (GC1) is presented in **Figure 2C**. This map elucidates the distribution of tissue architecture in GC, highlighting the spatial confinement of malignant-dominant CCZs (red) and fibroblast-dominant CCZs (green). Malignant-infiltrated CCZs (magenta) and fibroblast-infiltrated CCZs (yellow) are depicted as intermediate zones encircled by regions characterized by a high density of immune cells (immune-dominant CCZ, blue). Zones with normal epithelium (epithelial-dominant CCZ, green) appear as isolated regions within the tissue. Spatial maps for other GC cases are provided in **Supplementary Fig. 3**. The other immunogenic GCs, specifically GC2 and GC8, show CCZ distribution patterns similar to GC1, marked by a pronounced perimeter of immune cells around malignant and fibroblast-infiltrated zones. Epithelial GCs (GC3, GC5, and GC7; illustrated in **Supplementary Fig. 3**) predominantly feature malignant-dominant and malignant-infiltrated CCZs, with fibroblast-infiltrated zones immediately adjacent to the malignant regions. Interestingly, immune-dominant CCZs, though reduced, still serve as boundaries between malignant and fibroblast-associated areas in epithelial GCs. In fibrotic GCs (GC4, GC6, and GC9; **Supplementary Fig. 3**), fibroblast-related CCZs prevail, with a notable interaction between malignant cells and stromal cells within the malignant-infiltrated zones. Immune cells, albeit sparse, are found co-localized with stromal cells. Unlike other subtypes, fibrotic GCs exhibit a complete absence of immune-dominant CCZs. Therefore, the spatially architecture of GC is largely governed by cellular balance between malignant cells and fibroblasts along with the delineating presence of immune cells with their respective CCZs. This relationship leads to a marked reduction in immune demarcation in regions predominantly composed of malignant cells or fibroblasts, with the phenomenon being especially pronounced in fibrotic GCs.

**Figure 2.**
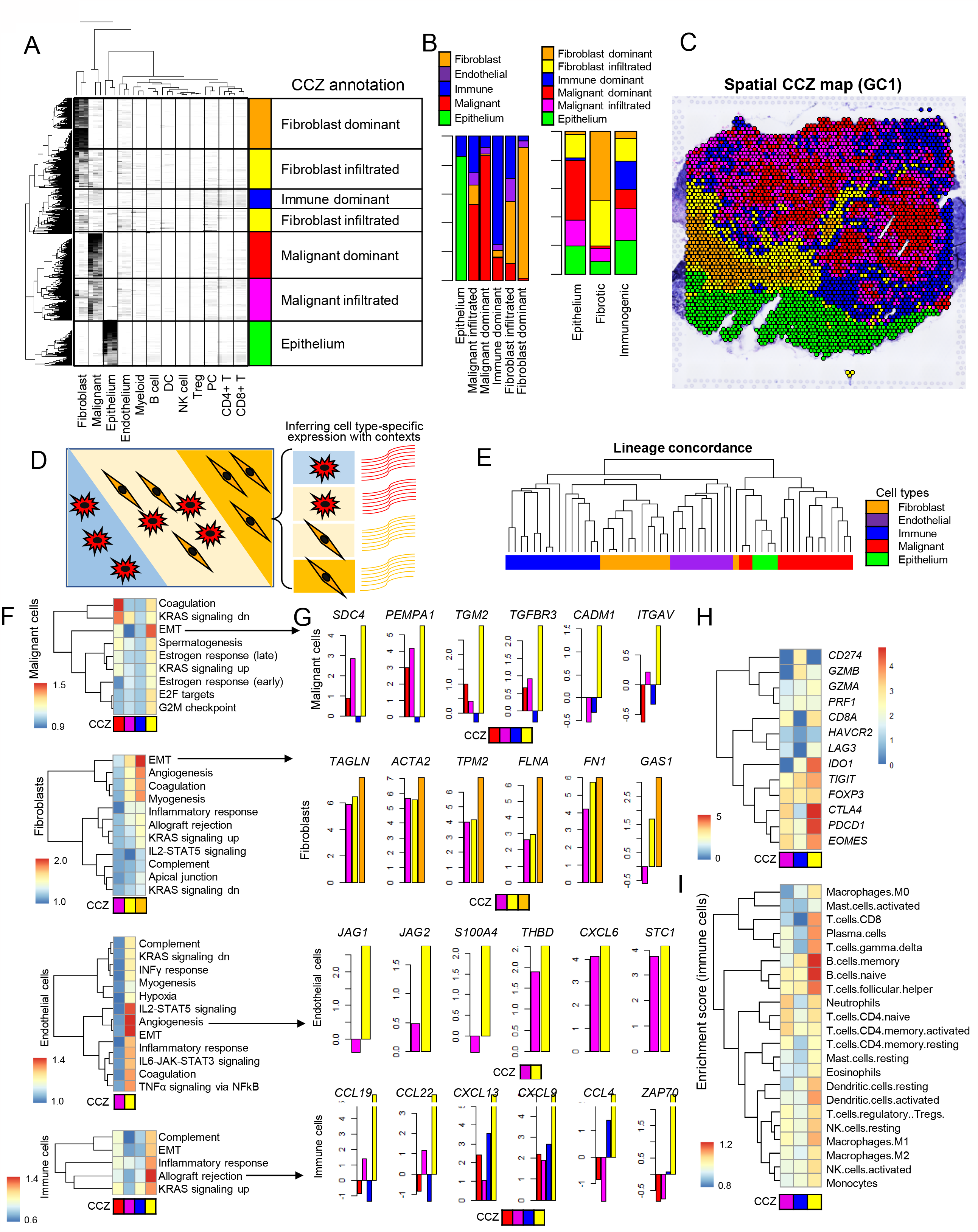
Cellular composite zone (CCZ) and their associated transcriptional dynamics of major TME cell types. (A) The total GC spots obtained across 9 cases (*n* = 29,808) are shown for their cellular abundance, which were subject to hierarchical clustering to segregate them into six CCZ (cellular composite zone) categories. Six CCZs were annotated with respect to the level of TME infiltration and major cell types. (B) Rhe distribution of cellular subtypes within each CCZ and their proportions in the three identified GC subtypes. (C) The spatial representation of CCZ annotations for case GC1. (D) A schematic representation of transcriptional dynamics inference in relation to CCZs, with examples of malignant cells and fibroblasts either isolated (left and right) or intermingled (middle) in three CCZs (left, middle and right). This led to inference of the cell type-specific expression patterns linked to different CCZs. (E) A high level of lineage concordance was observed for inferred cell type-specific expression across CCZs. (F) Functional enrichment analysis for four cell types (malignant cells, fibroblasts, endothelial, and immune cells) across different CCZs, with significant enrichment (FDR < 0.1) displayed in individual heatmaps. (G) For the selected functions (e.g., EMT in malignant cells), the genes whose expression changes across CCZs are shown. (H) and (I) show the expression of selected immune-related genes in immune cells and the CIBERSORT-estimated abundance of 22 immune cell types across CCZs, respectively.

Malignant-dominant and malignant-infiltrated CCZs correspond to histologically defined tumor cores and invasive margins. It is postulated that malignant cells within these distinct histological areas exhibit unique gene expression profiles, a hypothesis that extends to other cell types such as fibroblasts (22,23). Investigating the variations in expression between these cells can shed light on the transcriptional behavior of malignant and other cells in response to TME infiltrations. Considering that spot-level expression profiles encompass a mixture of various cell types, we employed deconvolution techniques to infer cell type-specific expression data for five primary cell types: malignant cells, normal epithelium, fibroblasts, endothelial cells, and aggregated immune cells. **Figure 2D** presents a schematic that visualizes the deconvolution process with reference to different CCZs. In this example, two cell types— malignant cells and fibroblasts—are shown in distinct spatial contexts: isolated regions (depicted on the left for malignant cells and on the right for fibroblasts) and areas of mutual infiltration (central region). The cell type-specific expression can be inferred in these defined regions (see *Methods*), allowing for a comparison that reveals the transcriptional dynamics of individual cell types with respect to TME contexts. For example, the expression profiles of malignant cells inferred from malignant-dominant regions are compared to those of malignant-infiltrated regions allowing for the identification of transcriptional dynamics of malignant cells influenced by fibroblast infiltration. Cell type-specific expression of five major cell types (malignant cells, fibroblasts, endothelial cells, and aggregated immune cells) were inferred across CCZs with substantial proportion of the cells of interests ensured (>5%). The inferred expression profiles were largely segregated in accordance with their cell lineages, confirming the accuracy of our cell type-specific expression profiling (**Fig. 2E**). Gene set enrichment analyses on four major cell types excluding normal epithelial cells identified key molecular functions as shown in heatmaps (significant enrichment of FDR < 0.1, **Fig. 2F**). The GC subtype-based functional enrichment analyses are shown in **Supplementary Fig. 4**. Malignant cells in malignant-dominant CCZs showed gene upregulation associated with ‘coagulation’. In contrast, malignant cells in fibroblast-infiltrated CCZs exhibited increased expression of genes linked to epithelial-to-mesenchymal transition (EMT). EMT, angiogenesis, and allograft rejection functions were notably upregulated in malignant cells, fibroblasts, endothelial cells, and immune cells within fibroblast-infiltrated CCZs, highlighting these as active zones in the GC tumor microenvironment (TME). For selected molecular functions (indicated by arrows in **Fig. 2F**), the expression level of pathway genes are shown across CCZs (**Fig. 2G**). For example, *SDC4, PEMPA1, TMG2, TGFBR3, CADM1*, and *ITGAV* as members of EMT signaling, showed transcriptional up-regulation in malignant cells residing in fibroblast-infiltrated CCZs and are responsible for the EMT activation of malignant cells in the milieu of highly infiltration of fibroblasts. In fibroblast-infiltrated and -dominant CCZs, the up-regulation of known cancer-associated fibroblasts (CAF) activation genes such as *TAGLN, ACTA2, TPM2*, and *FN1*, were observed for fibroblasts suggesting that the fibroblasts acquire the activated CAF properties in the fibroblast-enriched TME. For immune cells, *CCL19, CCL22, CXCL13, CXCL9, CCL4*, and *ZAP70* were transcriptionally up-regulated with the infiltration of fibroblasts. The transcriptional levels of immune-related genes, including various immune exhaustion markers like *IDO1, CTLA4, PDCD1*, and *EOMES*, were also elevated in immune cells in fibroblast-infiltrated CCZs (**Fig. 2H**). The analysis of 22 immune cell abundance also revealed a higher prevalence of B cells and myeloid cells (dendritic cells and M1 macrophages) in these CCZs, suggesting that the fibroblast infiltration reconstitute the immune cell compositions in GC TME (**Fig. 2I**).

### A landscape of intercellular crosstalk and functional consequences

While cell type-specific analysis with respect to CCZ underlines the transcriptional dynamics of individual cell types in relation to TME, it remains challenging to identify which cell types are responsible for such functional consequences of cellular interactions as regulators. To unravel the cellular interactions and their genetic mediators within the TME, we devised a method for inferring the crosstalk between ‘regulator’ and ‘target’ cell types and their subsequent functional outcomes, depicted in **Figure 3A**. Our strategy begins with establishing per-spot cell type-specific expression profile where the expression of five major cell types are inferred from the individual spots (as jointly analyzed with their matching 1^st^ and 2^nd^ neighbors to yield 19 spots). To infer the expression of five cell types, 20 spots are required (ref and **Supplementary Fig. 5**). Subsequently, we conducted GSEA on the inferred expression profiles of the ‘target’ cells to derive their functional activity scores. By examining the correlation between these functional activity profiles of target cells and the cellular abundance of ‘regulator’ cells, we could infer the influence of regulator cell infiltration on the function of target cells. The substantial level of correlation indicates that the infiltration of regulator cells has a functional impact on target cells, illuminating the cell-to-cell functional relationship. The comparison was performed in a pairwise manner for four major cell types (malignant cells, fibroblasts, endothelial cells and immune cells, excluding normal epithelium) yielding a landscape of functional crosstalk illustrated as a comprehensive heatmap (**Fig. 3B**).

**Figure 3.**
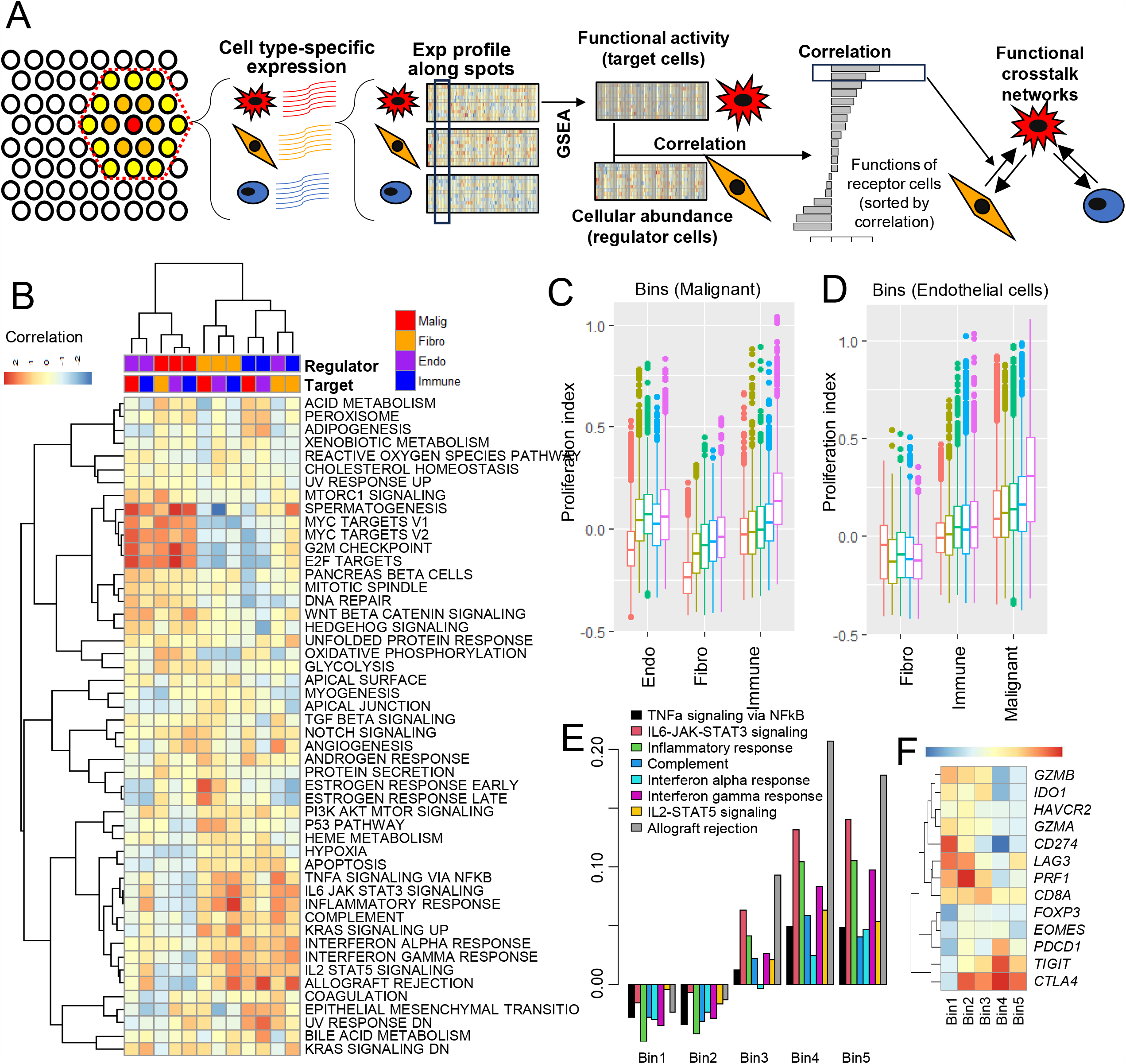
Landscape of cellular crosstalk in GC TME and their functional consequences. (A) Schematics are shown for the cell type-specific expression inference at spot levels and the building of functional crosstalk. For each spot, adjacent 18 spots (6 N1 and 12 N2 spots) were combined to infer the cell type-specific expression of major cell types. The deconvoluted expression of cell types were combined across spots and further subject to gene set enrichment analysis to obtain the functional scores of target cells. Across spots, the functional score profiles (target cells) were correlated with the cellular abundance of regulator cells. The high level of correlation indicates the functional relationship between target and receptors cells. (B) The ligand and target cell types (four) were compared in a pairwise manner. A heatmap represents the level of correlation for individual functional terms and each column corresponds to the regulator and target cells (top panels). For example, the leftmost column represents the endothelial cells as regulator cells may have impact on malignant cells as target cells leading to up-regulation of MYC targets in target cells. (C) Spots were binned in order of the infiltrating levels of malignant cells. The proliferation index was estimated using the genes associated with cellular proliferation, demonstrating the overall increase of proliferation index of major TME cells in GC with the infiltration of malignant cells. (D) Similarly shown for endothelial cells. (E) Across fibroblast-bins, the 8 immune functions activity was estimated for immune cell-specific expression. (F) Similarly, the level of immune related genes are shown across fibroblast bins.

The major functional changes in GC TME were the upregulation of genes associated with cell proliferation such as those of G2M checkpoint, MYC targets and E2F targets. This upregulation coincides with the infiltration of malignant and endothelial cells as regulator cells. This suggests that the cells in GC TME adapt by enhancing cellular proliferations in response to endothelial and malignant cell infiltration. To support the observation, we estimated proliferation index using the genes representing cell proliferation. The cell type-specific proliferation index were compared with respect to the level of infiltration of malignant cells (**Fig. 3C**) and endothelial cells (**Fig. 3D**) by categorizing spots into quintiles (bin1 to bin5, based on the level of malignant cell or endothelial cells). Notably, there was a consistent upward trend in the proliferation index correlating with these infiltrations, except in the context of the interaction between fibroblasts and endothelial cells. This trend underlines the potential roles of malignant and endothelial cells play within the GC TME. We also observed that the upregulation of various immune-related functions presented by 8 hallmark gene sets, were aligning with the infiltration of fibroblasts and immune cells as regulator cells. The 8 immune-related gene sets were further distinguished as those representing a general innate and adaptive immune response—such as responses to IFNα and IFNγ, IL2-STAT5 signaling, and allograft rejection, that are associated with the infiltration of immune cells and those relatively associated with fibroblasts’ regulatory influence over malignant and immune cells. This latter group includes pathways includes inflammation-related immune responses like TNFα signaling through NFκB, IL6-JAK-STAT3 signaling, the inflammatory response, and complement activation. Of interests, the infiltration of endothelial cells upregulated these classes of immune response instead of cellular proliferation of fibroblasts. We next focus on the impact of fibroblast infiltration on immune cells in terms of eight immune-related functions (**Fig. 3E**). In five bins with respect to fibroblast infiltration (bin1 and bin5 for those with the lowest and highest infiltration of fibroblasts), the scores of 8 immune-related gene sets showed a consistently increasing pattern including two functions of IL6-JAK-STAT3 and allograft rejection (**Fig. 3E**). We also examined the expression level of individual immune-related genes along the bins of fibroblast infiltration (**Fig. 3F**). While cytotoxic genes such as *GZMA* and *GZMB* were up-regulated in spots of low level of fibroblast infiltration, immune checkpoints including *PDCD1, CTLA4, TIGIT* were highly up-regulated along the fibroblast infiltrations suggesting that these immune checkpoints might be responsible for the immune cell dysfunctions regulated by the interaction with fibroblasts.

### *CCL2+* cancer-associated fibroblasts regulate the JAK-STAT3 signaling of macrophages

Among the immune functions of immune cells potentially modulated by fibroblasts (**Fig. 3E**), we focus on the ‘IL6-JAK-STAT3’ as related with inflammatory pro-cancer immune response in GC TME (24). We utilized NicheNet analysis (25) aiming to identify the key ligands in regulator cells (fibroblasts) and their matching receptor and target genes in receptor cells (immune cells). **Figure 4A** demonstrates the ligands expressed in fibroblasts as well as target and receptor genes expressed in immune cells. Among the ligands expressed in fibroblasts, *CCL2* emerged as a master regulator demonstrating extensive connection with targets gene belonging to IL6-JAK-STAT3 signaling (e.g., *LTB, SOCS1, SOCS3, STAT3, STAT3*, and *TGFB1*) and also the highest level of ligand activity as estimated by NicheNet analysis. Additional regulators expressed by fibroblasts include known CAF markers such as *TIMP-1* and *FN* (26).

**Figure 4.**
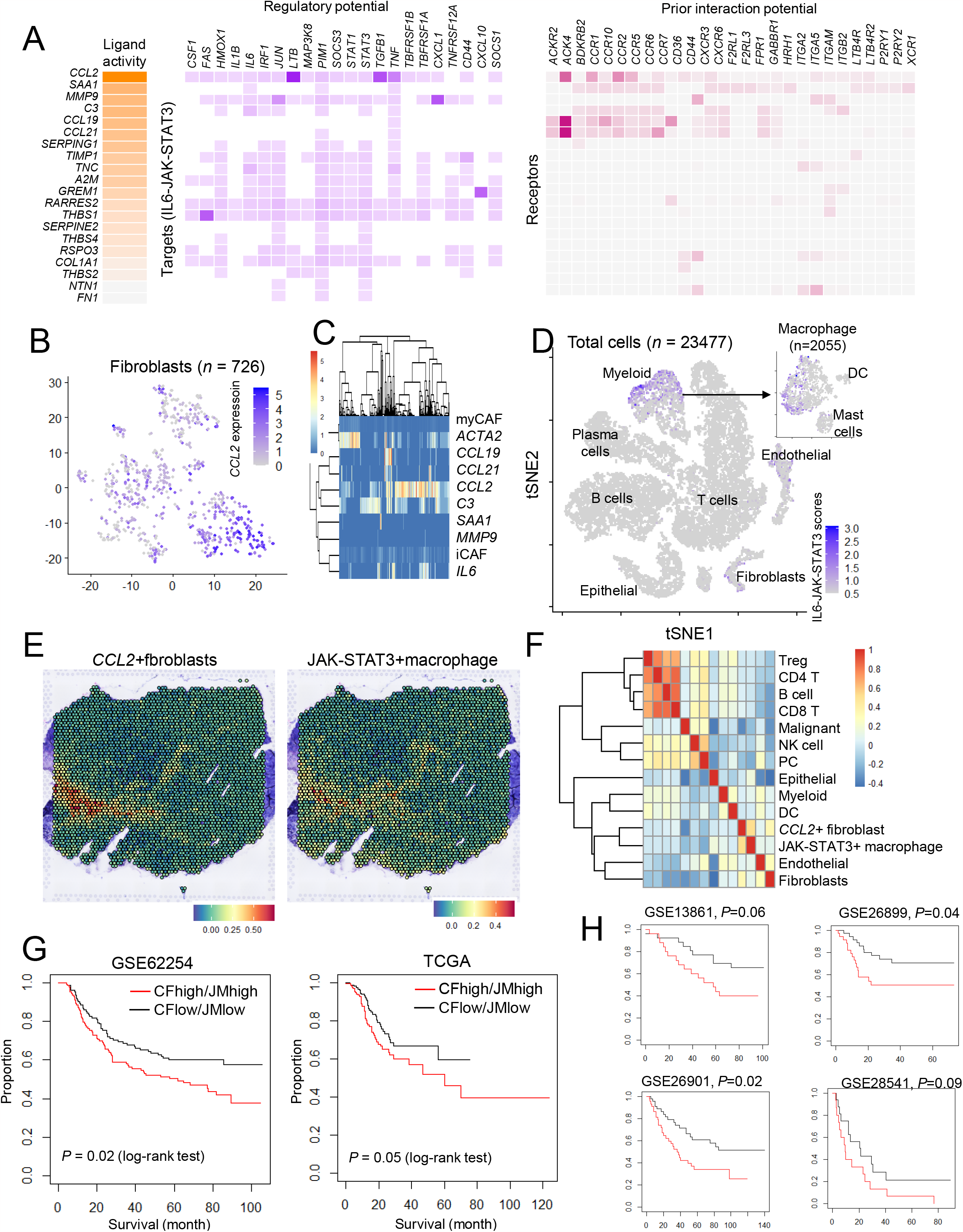
CCL2+fibroblasts and JAK-STAT3+macrophages with spatial colocalization and clinical outcomes. (A) NicheNet analysis reveals the top ligands associated with the cellular interactions between fibroblasts and immune cells, highlighting *CCL2* with top ligand activity. (B) tSNE plot showing distribution of CCL2 expression across 716 fibroblasts in scRNA-seq data. (C) Heatmap shows the level of expression of ligands including *CCL2* and scores of iCAF and myCAF. (D) A total GC TME cells (n = 23477) are shown in a tSNE plot and IL6-JAK-STAT3 signature scores are limited in myeloid cells. The inlet shows the myeloid cells among which IL6-JAK-STAT3 signature scores are also limited in macrophages. (E) In a GC1 spatial map, the spot scores representing *CCL2+*fibroblast and JAK-STAT3+macrophage show a concordant pattern suggestive of co-localization of two cell types. (F) Heatmap shows the correlation level measured across all the spots for the cellular abundance and the scores of two cell types (*CCL2+*fibroblast and JAK-STAT3+macrophage). (G) For six public GC bulk-level transcriptome data, *CCL2+*fibroblast-high (CF-high) and JAK-STAT3+macrophage-high (JM-high) samples (shown as red) shows substantially worse clinical outcomes compared to *CCL2+*fibroblast-low (CF-low) and JAK-STAT3+macrophage (JM-low) samples (shown as black).

To further characterize the *CCL2* expression of fibroblasts in GC TME, we employed our previous scRNA-seq data that include 726 fibroblasts obtained from five GC specimens (16). In a tSNE plot representing 726 fibroblasts, the expression level of *CCL2* is widely distributed rather than localized (**Fig. 2B**). We also explored the correlation between CCL2 expression and other ligands, as well as the scores for inflammatory CAF (iCAF) and myofibroblastic CAF (myCAF), as illustrated (**Fig. 2C**). Fibroblasts expressing *CCL2* comprise a substantial proportion of fibroblasts in GC TME, being mutually exclusive to *ACTA2+* myCAF but encompassing *IL6+* iCAF suggesting that *CCL2+*fibroblasts share transcriptional similarity with *IL6+*iCAF but represent a more prevalent fibroblast subclass in GC TME. Thus, we distinguish *CCL2+*fibrobalsts from *CCL2-*fibroblasts using a cutoff based on *CCL2* expression (143 CCL2+ fibroblasts *vs*. 583 CCL2-fibroblasts). The tSNE plots for fibroblasts markers and additional information are shown (**Supplementary Fig. S6**). Next, we examined the JAK-STAT activity (here, signature scores of ‘IL6-JAK-STAT3’ Hallmark gene set) in macrophages. In total single cells in GC TME (*n* = 23,477), the JAK-STAT3 score is predominantly found in myeloid cells, particularly in macrophages, indicating that macrophages are the primary target cells for CCL2+ fibroblasts among the immune cell types (**Fig. 3D**). Since the JAK-STAT3 activity is noted in a subset of macrophages, we also examined the expression of 7 macrophage markers (27) along with score of JAK-STAT3 and M1/M2 macrophages (**Supplementary Fig. S7**). Consequently, we categorized macrophages based on the activation of JAK-STAT3 signaling, using a cutoff for the JAK-STAT3 score (251 with activation *vs*. 1804 without).

Subsequently, we identified the characteristic genes for *CCL2+* fibroblasts and JAK-STAT3+ macrophages using the CIBERSORT signature matrix functions, selecting 425 signature genes for each cell type (details provided in **Supplementary Table S3**). We then examined if the spatial-level signature scores of *CCL2+* fibroblasts and JAK-STAT3+ macrophages exhibited co-localization in our spatial data. In the case of GC1, a strong agreement was observed between the abundance of *CCL2+* fibroblasts and JAK-STAT3+ macrophages based on their signature scores (**Fig. 4E**). The correlation between the abundance of major cell types and signature across of two cell types examined across all the spots of nine cases, indicated a high degree of correlation between *CCL2+* fibroblasts and JAK-STAT3+ macrophages, largely clustered to stromal cells but forming a distinct cluster (**Fig. 4F**). The spatial-level colocalization of these two cell types for the other cases is illustrated in **Supplementary Fig. S8**. However, despite this overall agreement, some cases demonstrated a predominance of either *CCL2+* fibroblasts (for example, GC9) or JAK-STAT3+ macrophages (for instance, GC4).

We further explored whether the signature score of CCL2+fibroblasts and JAK-STAT3+macrophages are associated with clinical outcomes. For clinical evaluation, we obtained public bulk-level RNA-seq or microarray-based GC data with patient outcomes. For GSE62254 (ACRG) dataset (n = 300), we classified the patients with respect to the scores of CCL2+fibroblasts and JAK-STAT3+macrophages with median values (CCL2+fibroblasts high/low as “CF-high/-low” and JAK-STAT3+macrophage high/low as “JM-high/-low”), leading to four categories (CF-high/JM-high, CF-high/JM-low, CF-low/JM-high, and CF-low/JM-low). For ACRG, We observed that 75% of the cases are CF-high/JM-high or CF-low/JM-low due to the strong concordance between two signature scores, and we focus on the comparison of the two classes (CF-high/JM-high vs. CF-low/JM-low) demonstrating that CF-high/JM-high GC show significantly worse prognosis (overall survival, P = 0.02, log-rank test) (Fig. 4G). The survival plots and patient numbers are fully presented in Supplementary Fig. S9. Similarly, TCGA and other additional four GC cohort showed similar clinical outcome.For clinical evaluation, we obtained public bulk-level RNA-seq or microarray based GC data with patient outcomes. For GSE62254 (ACRG) dataset (*n* = 300), we classified the patients with respect to the scores of *CCL2+*fibroblasts and JAK-STAT3+macrophages with median values, obtaining four categories (*CCL2+*fibroblasts high/low as “CF-high/-low” and JAK-STAT3+macrophage high/low as “JM-high/-low” resulting in four categories of CF-high/JM-high, CF-high/JM-low, CF-low/JM-high, and CF-low/JM-low). We found that more than 80% of the cases fell into either the CF-high/JM-high or CF-low/JM-low categories (84%, 252 out of 300 cases for ACRG and 87.6%, 338 out of 386 cases for TCGA), reflecting a strong correlation between the two signature scores. Thus, we focused on comparison of two groups (CF-high/JM-high *vs*. CF-low/JM-low). The analysis revealed that patients in the CF-high/JM-high category exhibited significantly poorer prognosis in terms of overall survival (*P* = 0.02, log-rank test, **Fig. 4G**). Complete details of the survival plots and patient numbers are provided in **Supplementary Figure S9**. Similar trends in clinical outcomes were observed in the TCGA (*P* = 0.05, log-rank test) and four other GC cohorts (GSE13861, GSE16899, GSE16901, GSE18541 with P = 0.06, 0.04, 0.02, 0.09, respectively) (**Fig. 5G**).

## Discussion

In this study, we utilized barcode-based Visium technology (19) for analyzing TME in nine GC specimens. This approach allowed for the detailed examination of the GC TME by incorporating tissue architecture into the transcriptomic analysis, thus facilitating the mapping of cellular interactions and their functional interpretations in GC TME. Differing from scRNA-seq, this technology employs unique molecular identifiers (UMIs) linked to spatially distinct spots, typically containing 3-10 cells each. This presents a challenge in differentiating individual cells within these spots, which often comprise various cell types (28). To overcome these technological limitations, deconvolution methods have become essential for analyzing Visium-based spatial transcriptomics data with comprehensive reviews and benchmarks available in other studies (29,30). Using cell type-specific abundance data, our analysis has delineated the spatial architecture of three primary gastric cancer (GC) subtypes: ‘immunogenic’, ‘epithelial’, and ‘fibrotic’. These subtypes, identified by distinct cellular compositions, align with the histology-based Lauren’s classification, e.g., intestinal type GCs are mostly categorized as immunogenic or epithelial, whereas diffuse type GCs are typically fibrotic. Spatial information further provides detailed insights into the interactions between malignant and tumor microenvironment (TME) cells within GC tissue architecture. For instance, immunogenic GCs like GC1, characterized by a diverse immune cell population including T cells and plasma cells, indicate a robust immune-hot TME previously classified as ‘immune-inflamed’ in GC (31) or ‘immune-hot’ tumors in PanCancer (32). These tumors demonstrate notable immune cell infiltration around malignant cells, suggesting active tumor-immune system interactions. It is of note that CCZ analysis reveals that these immune cells are mostly observed as intermixed with other cell types rather than present solely in spot-resolution spatial dataset. A marked reduction in T and plasma cells, but a consistent presence of myeloid cells and B cells in epithelial GC subtypes as well as the extensive fibroblast infiltration in fibrotic GCs are unique GC subtype-specific TME features. The presence of various immune cells, despite a lack of T cells, indicates complex interactions within the TME that may affect disease progression and treatment response.

The abundance of cells can be also determined using scRNA-seq, but spatial data offers unique insights into the spatial arrangement in GC subtypes. Thus, we have identified spatial-level spot clusters, termed CCZs, which allow us to categorize regional spots in the GC TME into at least six distinct categories. To understand the transcriptional dynamics of these CCZs, we utilized a method of expression deconvolution specific to each cell type. Initial methods for deconvolving spatial transcriptome data primarily estimated the abundance of specific cell types in spots (30). However, some of these techniques also sought to discern the expression profiles of individual cell types within particular tissue environments aiming to deconvolute cell type-specific or signature expressions, starting from cell abundance data, a process called reference-free deconvolution (33,34). These methods have been applied in bulk-level RNA-seq (35) to create PanCancer-scale tumor ecotypes (36). Yet, in spatial transcriptomics, particularly in cancerous tissues, deconvoluting cell type-specific expressions is more complex and has mostly been tested in simpler, non-cancerous structures at the tissue-level transcriptome deconvolution (e.g., whole spots in given context) (33,34). To overcome the sparsity of the datasets and complexity of GC spatial transcriptomics data, we limited the number of cell types to be deconvoluted (e.g., five major cell types of malignant, normal epithelium, fibroblasts, endothelial and immune cells) (35). We also used CCZ-based spot annotations or regional neighbors as a unit for group optimization in expression inference (35).

Our analysis revealed that in the fibroblast-enriched GC TME, the major cell types exhibit up-regulation of their characteristic functions, such as EMT in malignant cells, angiogenesis in endothelial cells, and allograft rejection in immune cells. This suggests that that CAF-enriched TME supports the GC promoting environments for major GC cellular components including malignant cells. In addition, while EMT is also heightened in fibroblasts within fibroblast-enriched CCZs, the EMT-related genes in fibroblasts (mainly CAF activation markers) differ from those in malignant cells (mainly TGFβ signaling genes). This suggests that a fibroblast-enriched TME may facilitate CAF-mediated TGFβ signaling activation in malignant cells and CAF activation in fibroblasts (37). Next, we exploited neighboring 18 spots (i.e., 6 N1 neighbors and 12 N2 neighbors) to given spots as ‘groups’ to deconvolute the spot-level gene expression. This strategy enabled us to build a landscape of cellular crosstalk between individual cell type revealing that endothelial and malignant cells infiltrating the TME might promote tumor growth by upregulating genes linked to cell proliferation. This finding is consistent with previous observation where TME respond to proliferating malignant cells, e.g., the promotion of angiogenesis to restore oxygen and the transformation of tumor infiltrating immune cells toward pro- or anti-tumorigenic functions (38). It is also possible that the increased endothelial cell infiltration leads to defective angiogenesis and the resulting hypoxic conditions may be associated with overall cellular proliferations of the TME cells (39).

Of interests, fibroblasts and immune cell infiltration into TME seem to enhance immune-related functions across different cell types. Among the functions, we highlighted IL6-JAK-STAT3 and inflammatory signaling in immune cells that may be regulated by tumor infiltrating fibroblasts. Of note, the signaling is restricted in macrophages among immune cells, thus raising a hypothesis that *CCL2+*fibroblasts trigger pro-cancer inflammatory response of macrophages in GC TME. JAK-STAT3 signaling has been well recognized for their roles in protumor signaling and studies have suggested they also play roles in the interactions between fibroblasts and macrophages (40). We further define that *CCL2+*fibroblasts and JAK-STAT3+macrophages may present as separate cell subpopulations. Their co-localization in spatial data further suggests that the interaction between two cell types implicates the presence of the inflammatory pro-cancer immune response within the GC TME. The study also extends its findings to clinical cohorts, demonstrating that the co-infiltration of *CCL2+*fibroblasts and JAK-STAT+macrophages is associated with unfavorable clinical outcomes in GC patients. This suggests that targeting these specific cellular interactions and modulating TME, could be a promising avenue for therapeutic interventions and personalized treatment strategies.

In summary, this study’s results provide valuable insights into the spatial and cellular heterogeneity of GC, shedding light on the complex interplay of various cell types within the TME. These findings have the potential to inform the development of novel therapeutic approaches and contribute to a better understanding of GC progression and prognosis. However, further research and clinical validation are needed to translate these findings into actionable strategies for GC patients.

## Methods

### Patient enrollment

Nine patients, diagnosed with gastric cancer (GC), were enrolled in this study after providing informed consent. The study received approval from the institutional review board (AJOUIRB-SM-2023-326). Detailed clinicopathological data about these patients can be found in **Supplementary Table 1**. Tumor specimens were collected during the surgical resection of the tumors. Surgical specimens were incubated in OCT (Optimal Cutting Temperature, 10X Visium) compound at a controlled temperature of 4°C for 30 minutes. Then, the specimens were frozen in OCT and stored at -80°C to maintain their integrity until they were prepared for further tissue analysis.

### Tissue preparation

Tissues were prepared as 10μm sections and placed onto Visium Gateway gene expression slides for staining and library preparation. Prepared tissue sections were stained with Hematoxylin and Eosin (H&E). Board-certified pathologists then examined H&E stained slides to identify areas of interests for transcriptome capture. The tissue sections underwent initial pre-permeabilization steps also a washing process. Then, tissue sections were permeabilized to capture RNA molecules onto barcoded spots on the slides. After permeabilization, we proceeded with library preparation, according to the manufacturer’s protocol for reverse transcription, to prepare the tissues for detailed transcriptomic analysis.

### Sequencing and data processing

The prepared libraries were sequenced using the Illumina NovaSeq (Illumina NovaSeq 6000) platform. Nine specimens yielded between 157 to 214 million raw reads with details on the sequencing information of each specimen available in **Supplementary Table 2**. We processed the raw sequencing data with the SpaceRanger software (version 1.7.0 from 10X Genomics) utilizing the human genome assembly GRCh38 (version 96) and its corresponding GENCODE annotation file (version 25) as the reference. In the preprocessing phase, we demultiplexed the base call (bcl) files, followed by the trimming of fastq files using Cutadapt, to eliminate any non-internal 5’ adaptor sequences and polyA homopolymers from the 3’ end of the reads. The SpaceRanger pipeline processed the trimmed data, mapping it to the GRCH38 as recommended by Manufacturer protocols (https://support.10xgenomics.com/single-cell-gene-expression/software/pipelines/latest/).

### Analysis of mixed cell populations with deconvolution

To address the challenge of mixed cell populations within the regional spots of transcriptome data, we applied various deconvolution methods to estimate the proportions of different cell types in GC TME. These methods included SPOTlight (41) and cell2location (42) as well as built-in functions of the Seurat software (43). For reference, we used single-cell data of five GC from our previous studies with annotations for 11 major cell types (16). To select the best-performing deconvolution method, we used histological annotations provided by a pathologist, which categorized the regions into malignant, normal, stromal, and immune areas. Comparisons were made between the histology-based and deconvolution-based spot annotations (i.e., those labeled based on the predominant cell types identified by deconvolution). By comparing three deconvolution algorithms (Seurat, SPOTlight and cell2location) in terms of their concordance with histological annotations by pathologists, Seurat showed balanced performance in terms of concordance as shown in **Supplementary Fig. 10**. Thus, Seurat-based deconvolution results were employed for the subsequent analyses. Based on the deconvolution results, we categorized the examined cases into three GC subtypes with respect to the cellular abundances of 12 cell types using hierarchical clustering (normal and malignant epithelial cells, fibroblasts, endothelial cells along with 8 immune cell types). According to the major cell types, the subtypes were annotated as epithelial, immunogenic and fibrotic GCs with the predominance of malignant cells, immune cells and fibroblasts, respectively. The spatial visualizations were made using Seurat software (43). The spatial relationship across the cell types were also examined by correlation coefficients of spot-level cellular abundance in given case.

### Cellular composite zones and cell type-specific expression

The units of spatial transcriptome data, the spots were organized in a centered rectangular lattice, ensuring that every spot was evenly bordered by six adjacent spots, referred to as ‘1st tier neighbors or N1 spots’. Surrounding these, the ‘2nd tier neighbors or N2 spots’ comprised 12 spots adjacent to the 1st tier neighbors. The abundance of 12 cell types were collected for individual spots as well as for 1^st^ and 2^nd^ neighboring spots (mean abundance for neighboring spots). The spot-level cellular abundance profile was subject to hierarchical clustering yielding six categories and annotated as CCZ (cellular composite zone). Six CCZs were annotated according to the major cell types and the level of TME infiltrations. Cell type-specific expression of each CCZ have been inferred by *nnls* (non-negative least square) based gene expression devolution. For deconvolution, we used cellular abundance for five cell types (normal and malignant epithelial cells, fibroblasts, endothelial cells, and immune cells) where we aggregated the cellular abundance of eight immune cell subtypes as that of immune cells. The cellular abundance was used to derive the cell type-specific expression of five cell types as described previously (35). The lineage concordance of cell type-specific expression was evaluated by hierarchical clustering. Cell type-specific expression obtained across CCZ were further subject to functional enrichment analysis (44) using Hallmark gene sets (https://www.gsea-msigdb.org/gsea/msigdb) and GSVA R packages with ssgsea options (45). The imputation of abundance for major 22 immune cell types were done across CCZ- and cell type-specific immune cell expression with CIBERSORT and LM22 signatures (46).

### Crosstalk landscape

To build a landscape of cellular crosstalk in GC TME, we performed deconvolution of cell type-specific expression for five major cell types across regional spots. For robust deconvolution, we collected 19 spots (spots of interests along with 6 and 12 1^st^ and 2^nd^ neighbors together) per each spot. We used nnls to deconvolute the cell type-specific expression and further used GSEA to obtain the spot-level functional scores corresponding to 50 terms of Hallmar gene sets. Interaction between two cell types (regulator cells and target cells) were estimated by a level of correlation between the spot-level functional scores of cell type A (target cell) were correlated with the abundance of cell type B (regulator cells) across the spots. The substantial level of correlated were translated into the functional relationship between cell type A and B as targets and regulators, respectively. We performed the analyses for the pairwise relationship across four major cell types (malignant cells, fibroblasts, endothelial cells and immune cells). To assess the level of cellular proliferation, we employed 157 genes associated with cell cycle and proliferation (47). The proliferation index was calculated as the mean of expression for genes. The proliferation index of individual spots were grouped into five bins with respect to the level of malignant and endothelial cell infiltration (i.e., the cellular abundance of the corresponding cell types in spots). To evaluate the immune response of immune cells and fibroblasts, we also employed 8 immune-related gene sets from Hallmark datasets and immune scores were similarly calculated with visualization across five bins with respect to the level of fibroblasts abundance.

### Subtyping of cell and colocalization

We utilized NicheNet analysis to identify interactions between fibroblasts (as the “regulator” cells) and immune cells (as the “target” cells) through ligand-receptor pairs and target genes. For NicheNet analysis, the genes expressed in fibroblasts and immune cells were used as input with default sets of ligand-receptor pairs and ligand-target regulatory scores provided with NicheNet (25). To refine specific cell subtypes involved in these interactions, we referred to our prior single-cell RNA-seq data from five gastric cancer samples (16). We focused on 726 fibroblasts, analyzing their expression of selected markers. To reveal the relationships between fibroblast markers including *CCL2* and other markers, we generated a heatmap with hierarchical clustering. For the study of inflammatory and myogenic cancer-associated fibroblasts (iCAF and myCAF, respectively), we looked at genes linked to these two subtypes, with *IL6* and *ACTA2* serving as key markers for iCAFs and myCAFs, respectively. Genes representing iCAF and myCAF were also obtained from a previous literature (48). For macrophages, we began with exclude mast cells and dendritic cells from myeloid population in GC TME. For macrophage (*n* = 2055), the ‘JAK-STAT” scores represented the average expression of genes belonging to the functional gene set of IL6-JAK-STAT3, were calculated. JAK-STAT3 scores of macrophages were compared with 7 macrophage markers previously proposed (*FCN1, NLRP3, PLTP, CXCL12, C1QC, SPP1*, and *MKI67*) (27) as well as M1 and M2 macrophage gene signatures (49). The marker genes representing *CCL2+*fibroblasts and JAK-STAT3+macrophages were identified using CIBERSORT signature matrix functions (35). The selected signature genes and their mean expression levels as scores representing the abundance of *CCL2+*fibroblast and JAK-STAT3+macrophage in spatial transcriptomic data spots and in bulk data for clinical association studies.

### Survival analyses

To validation the clinical relevance of the infiltration of *CCL2+*fibroblasts and JAK-STAT3+macrophages in GC TME, we downloaded the bulk-level transcriptome data of TCGA consortium (2), ACRG consortium (GSE62254) (10) as well as GSE13861 (50), GSE26899 (51), GSE26901 (51), and GSE28541 (51). RNA-seq and microarray-based expression profiles were used as obtained from GEO database and the scores based on mean expression of marker genes for *CCL2+*fibroblasts and JAK-STAT3+macrophages were calculated. We divide the patients by the median of individual scores to categorize the patients into four categories of CCL2+fibroblast-high and JAK-STAT3+macrophage-high (CF-high/JK-high) as well as CF-high/JM-low, CF-low/JM-high and CF-low/JM-low. The survival analysis was done for patients’ overall survival using log-rank tests with Kaplan-Meyer curves.

## Data availability

The sequencing data has been uploaded to GEO (gene expression omnibus) with accession ID (GSE251950).

## Acknowledgements

We appreciate the support of this research by Basic Medical Science Facilitation Program through the Catholic Medical Center of the Catholic University of Korea funded by the Catholic Education Foundation

## References

1. Yeoh KG, Tan P. Mapping the genomic diaspora of gastric cancer. Nat Rev Cancer 2021 doi 10.1038/s41568-021-00412-7.

2. Cancer Genome Atlas Research N. Comprehensive molecular characterization of gastric adenocarcinoma. Nature 2014;513(7517):202–9 doi 10.1038/nature13480.

3. Zang ZJ, Cutcutache I, Poon SL, Zhang SL, McPherson JR, Tao J, et al. Exome sequencing of gastric adenocarcinoma identifies recurrent somatic mutations in cell adhesion and chromatin remodeling genes. Nat Genet 2012;44(5):570–4 doi 10.1038/ng.2246.

4. Wang K, Yuen ST, Xu J, Lee SP, Yan HH, Shi ST, et al. Whole-genome sequencing and comprehensive molecular profiling identify new driver mutations in gastric cancer. Nat Genet 2014;46(6):573–82 doi 10.1038/ng.2983.

5. Yoo HY, Sung MK, Lee SH, Kim S, Lee H, Park S, et al. A recurrent inactivating mutation in RHOA GTPase in angioimmunoblastic T cell lymphoma. Nat Genet 2014;46(4):371–5 doi 10.1038/ng.2916.

6. Nakamura Y, Kawazoe A, Lordick F, Janjigian YY, Shitara K. Biomarker-targeted therapies for advanced-stage gastric and gastro-oesophageal junction cancers: an emerging paradigm. Nat Rev Clin Oncol 2021;18(8):473–87 doi 10.1038/s41571-021-00492-2.

7. Bang YJ, Van Cutsem E, Feyereislova A, Chung HC, Shen L, Sawaki A, et al. Trastuzumab in combination with chemotherapy versus chemotherapy alone for treatment of HER2-positive advanced gastric or gastro-oesophageal junction cancer (ToGA): a phase 3, open-label, randomised controlled trial. Lancet 2010;376(9742):687–97 doi 10.1016/S0140-6736(10)61121-X.

8. Wang R, Dang M, Harada K, Han G, Wang F, Pool Pizzi M, et al. Single-cell dissection of intratumoral heterogeneity and lineage diversity in metastatic gastric adenocarcinoma. Nat Med 2021;27(1):141–51 doi 10.1038/s41591-020-1125-8.

9. Pectasides E, Stachler MD, Derks S, Liu Y, Maron S, Islam M, et al. Genomic Heterogeneity as a Barrier to Precision Medicine in Gastroesophageal Adenocarcinoma. Cancer Discov 2018;8(1):37–48 doi 10.1158/2159-8290.CD-17-0395.

10. Cristescu R, Lee J, Nebozhyn M, Kim KM, Ting JC, Wong SS, et al. Molecular analysis of gastric cancer identifies subtypes associated with distinct clinical outcomes. Nat Med 2015;21(5):449–56 doi 10.1038/nm.3850.

11. Yeoh KG, Tan P. Mapping the genomic diaspora of gastric cancer. Nat Rev Cancer 2022;22(2):71–84 doi 10.1038/s41568-021-00412-7.

12. Zhang P, Yang M, Zhang Y, Xiao S, Lai X, Tan A, et al. Dissecting the Single-Cell Transcriptome Network Underlying Gastric Premalignant Lesions and Early Gastric Cancer. Cell Rep 2019;27(6):1934–47 e5 doi 10.1016/j.celrep.2019.04.052.

13. Sathe A, Grimes SM, Lau BT, Chen J, Suarez C, Huang RJ, et al. Single-Cell Genomic Characterization Reveals the Cellular Reprogramming of the Gastric Tumor Microenvironment. Clin Cancer Res 2020;26(11):2640–53 doi 10.1158/1078-0432.CCR-19-3231.

14. Zhang M, Hu S, Min M, Ni Y, Lu Z, Sun X, et al. Dissecting transcriptional heterogeneity in primary gastric adenocarcinoma by single cell RNA sequencing. Gut 2021;70(3):464–75 doi 10.1136/gutjnl-2019-320368.

15. Kumar V, Ramnarayanan K, Sundar R, Padmanabhan N, Srivastava S, Koiwa M, et al. Single-Cell Atlas of Lineage States, Tumor Microenvironment, and Subtype-Specific Expression Programs in Gastric Cancer. Cancer Discov 2022;12(3):670–91 doi 10.1158/2159-8290.CD-21-0683.

16. Jeong HY, Ham IH, Lee SH, Ryu D, Son SY, Han SU, et al. Spatially Distinct Reprogramming of the Tumor Microenvironment Based On Tumor Invasion in Diffuse-Type Gastric Cancers. Clin Cancer Res 2021;27(23):6529–42 doi 10.1158/1078-0432.CCR-21-0792.

17. Asp M, Bergenstrahle J, Lundeberg J. Spatially Resolved Transcriptomes-Next Generation Tools for Tissue Exploration. Bioessays 2020;42(10):e1900221 doi 10.1002/bies.201900221.

18. Quail DF, Joyce JA. Microenvironmental regulation of tumor progression and metastasis. Nat Med 2013;19(11):1423–37 doi 10.1038/nm.3394.

19. Stahl PL, Salmen F, Vickovic S, Lundmark A, Navarro JF, Magnusson J, et al. Visualization and analysis of gene expression in tissue sections by spatial transcriptomics. Science 2016;353(6294):78–82 doi 10.1126/science.aaf2403.

20. Cho J, Chang YH, Heo YJ, Kim S, Kim NK, Park JO, et al. Four distinct immune microenvironment subtypes in gastric adenocarcinoma with special reference to microsatellite instability. ESMO Open 2018;3(3):e000326 doi 10.1136/esmoopen-2018-000326.

21. Ji AL, Rubin AJ, Thrane K, Jiang S, Reynolds DL, Meyers RM, et al. Multimodal Analysis of Composition and Spatial Architecture in Human Squamous Cell Carcinoma. Cell 2020;182(2):497–514 e22 doi 10.1016/j.cell.2020.05.039.

22. Arora R, Cao C, Kumar M, Sinha S, Chanda A, McNeil R, et al. Spatial transcriptomics reveals distinct and conserved tumor core and edge architectures that predict survival and targeted therapy response. Nat Commun 2023;14(1):5029 doi 10.1038/s41467-023-40271-4.

23. Xun Z, Ding X, Zhang Y, Zhang B, Lai S, Zou D, et al. Reconstruction of the tumor spatial microenvironment along the malignant-boundary-nonmalignant axis. Nat Commun2023;14(1):933 doi 10.1038/s41467-023-36560-7.

24. Owen KL, Brockwell NK, Parker BS. JAK-STAT Signaling: A Double-Edged Sword of Immune Regulation and Cancer Progression. Cancers (Basel) 2019;11(12) doi 10.3390/cancers11122002.

25. Browaeys R, Saelens W, Saeys Y. NicheNet: modeling intercellular communication by linking ligands to target genes. Nat Methods 2020;17(2):159–62 doi 10.1038/s41592-019-0667-5.

26. Zhang H, Yue X, Chen Z, Liu C, Wu W, Zhang N, et al. Define cancer-associated fibroblasts (CAFs) in the tumor microenvironment: new opportunities in cancer immunotherapy and advances in clinical trials. Mol Cancer 2023;22(1):159 doi 10.1186/s12943-023-01860-5.

27. Liu Y, Zhang Q, Xing B, Luo N, Gao R, Yu K, et al. Immune phenotypic linkage between colorectal cancer and liver metastasis. Cancer Cell 2022;40(4):424–37 e5 doi 10.1016/j.ccell.2022.02.013.

28. Williams CG, Lee HJ, Asatsuma T, Vento-Tormo R, Haque A. An introduction to spatial transcriptomics for biomedical research. Genome Med 2022;14(1):68 doi 10.1186/s13073-022-01075-1.

29. Li B, Zhang W, Guo C, Xu H, Li L, Fang M, et al. Benchmarking spatial and single-cell transcriptomics integration methods for transcript distribution prediction and cell type deconvolution. Nat Methods 2022;19(6):662–70 doi 10.1038/s41592-022-01480-9.

30. Li H, Zhou J, Li Z, Chen S, Liao X, Zhang B, et al. A comprehensive benchmarking with practical guidelines for cellular deconvolution of spatial transcriptomics. Nat Commun 2023;14(1):1 48 doi 10.1038/s41467-023-37168-7.

31. Chen Y, Sun Z, Chen W, Liu C, Chai R, Ding J, et al. The Immune Subtypes and Landscape of Gastric Cancer and to Predict Based on the Whole-Slide Images Using Deep Learning. Front Immunol 2021;12:685992 doi 10.3389/fimmu.2021.685992.

32. Bagaev A, Kotlov N, Nomie K, Svekolkin V, Gafurov A, Isaeva O, et al. Conserved pan-cancer microenvironment subtypes predict response to immunotherapy. Cancer Cell 2021;39(6):845–65 e7 doi 10.1016/j.ccell.2021.04.014.

33. Ma Y, Zhou X. Spatially informed cell-type deconvolution for spatial transcriptomics. Nat Biotechnol 2022;40(9):1349–59 doi 10.1038/s41587-022-01273-7.

34. Miller BF, Huang F, Atta L, Sahoo A, Fan J. Reference-free cell type deconvolution of multicellular pixel-resolution spatially resolved transcriptomics data. Nat Commun 2022;13(1):2339 doi 10.1038/s41467-022-30033-z.

35. Newman AM, Steen CB, Liu CL, Gentles AJ, Chaudhuri AA, Scherer F, et al. Determining cell type abundance and expression from bulk tissues with digital cytometry. Nat Biotechnol 2019;37(7):773–82 doi 10.1038/s41587-019-0114-2.

36. Margolin AA, Nemenman I, Basso K, Wiggins C, Stolovitzky G, Dalla Favera R, et al. ARACNE: an algorithm for the reconstruction of gene regulatory networks in a mammalian cellular context. BMC Bioinformatics 2006;7 Suppl 1:S7 doi 10.1186/1471-2105-7-S1-S7.

37. Yu Y, Xiao CH, Tan LD, Wang QS, Li XQ, Feng YM. Cancer-associated fibroblasts induce epithelial-mesenchymal transition of breast cancer cells through paracrine TGF-beta signalling. Br J Cancer 2014;110(3):724–32 doi 10.1038/bjc.2013.768.

38. Anderson NM, Simon MC. The tumor microenvironment. Curr Biol 2020;30(16):R921–R5 doi 10.1016/j.cub.2020.06.081.

39. Feitelson MA, Arzumanyan A, Kulathinal RJ, Blain SW, Holcombe RF, Mahajna J, et al. Sustained proliferation in cancer: Mechanisms and novel therapeutic targets. Semin Cancer Biol 2015;35 Suppl(Suppl):S25–S54 doi 10.1016/j.semcancer.2015.02.006.

40. Witherel CE, Abebayehu D, Barker TH, Spiller KL. Macrophage and Fibroblast Interactions in Biomaterial-Mediated Fibrosis. Adv Healthc Mater 2019;8(4):e1801451 doi 10.1002/adhm.201801451.

41. Elosua-Bayes M, Nieto P, Mereu E, Gut I, Heyn H. SPOTlight: seeded NMF regression to deconvolute spatial transcriptomics spots with single-cell transcriptomes. Nucleic Acids Res 2021;49(9):e50 doi 10.1093/nar/gkab043.

42. Kleshchevnikov V, Shmatko A, Dann E, Aivazidis A, King HW, Li T, et al. Cell2location maps fine-grained cell types in spatial transcriptomics. Nat Biotechnol 2022;40(5):661–71 doi 10.1038/s41587-021-01139-4.

43. Hao Y, Stuart T, Kowalski MH, Choudhary S, Hoffman P, Hartman A, et al. Dictionary learning for integrative, multimodal and scalable single-cell analysis. Nat Biotechnol 2023 doi 10.1038/s41587-023-01767-y.

44. Subramanian A, Tamayo P, Mootha VK, Mukherjee S, Ebert BL, Gillette MA, et al. Gene set enrichment analysis: a knowledge-based approach for interpreting genome-wide expression profiles. Proc Natl Acad Sci U S A 2005;102(43):15545–50 doi 10.1073/pnas.0506580102.

45. Hanzelmann S, Castelo R, Guinney J. GSVA: gene set variation analysis for microarray and RNA-seq data. BMC Bioinformatics 2013;14:7 doi 10.1186/1471-2105-14-7.

46. Newman AM, Liu CL, Green MR, Gentles AJ, Feng W, Xu Y, et al. Robust enumeration of cell subsets from tissue expression profiles. Nat Methods 2015;12(5):453–7 doi 10.1038/nmeth.3337.

47. Locard-Paulet M, Palasca O, Jensen LJ. Identifying the genes impacted by cell proliferation in proteomics and transcriptomics studies. PLoS Comput Biol 2022;18(10):e1010604 doi 10.1371/journal.pcbi.1010604.

48. Ohlund D, Handly-Santana A, Biffi G, Elyada E, Almeida AS, Ponz-Sarvise M, et al. Distinct populations of inflammatory fibroblasts and myofibroblasts in pancreatic cancer. J Exp Med 2017;214(3):579–96 doi 10.1084/jem.20162024.

49. Najafi M, Hashemi Goradel N, Farhood B, Salehi E, Nashtaei MS, Khanlarkhani N, et al. Macrophage polarity in cancer: A review. J Cell Biochem 2019;120(3):2756–65 doi 10.1002/jcb.27646.

50. Cho JY, Lim JY, Cheong JH, Park YY, Yoon SL, Kim SM, et al. Gene expression signature-based prognostic risk score in gastric cancer. Clin Cancer Res 2011;17(7):1850–7 doi 10.1158/1078-0432.CCR-10-2180.

51. Oh SC, Sohn BH, Cheong JH, Kim SB, Lee JE, Park KC, et al. Clinical and genomic landscape of gastric cancer with a mesenchymal phenotype. Nat Commun 2018;9(1):1777 doi 10.1038/s41467-018-04179-8.

