## Supplementary Figs for "Spatial dissection of cellular heterogeneity and functional crosstalk in the tumor microenvironments of gastric cancers"

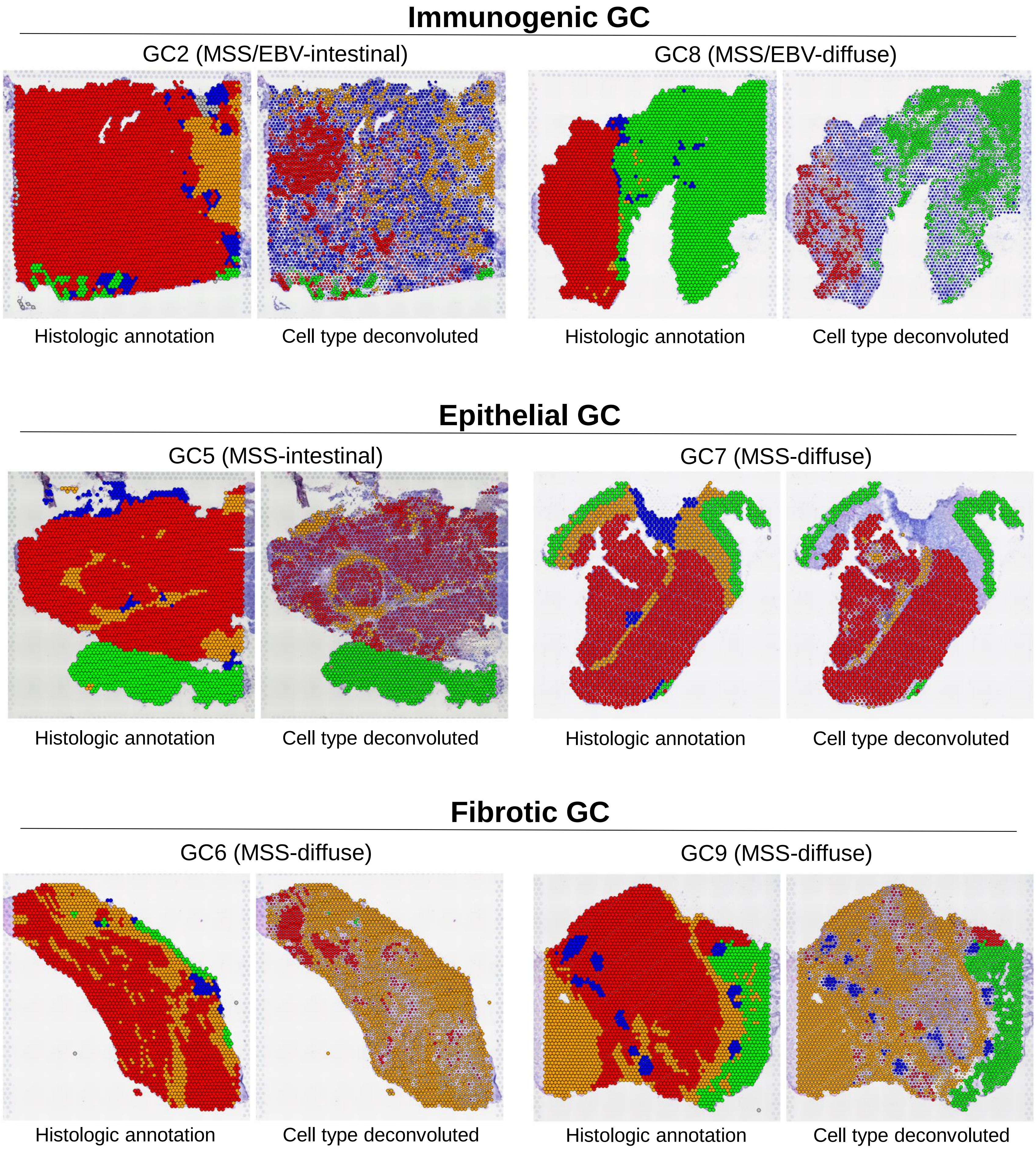


**Supplementary Fig. 1. Spatial cellular map of three GC subtypes.** For the remaining cases (two cases per GC subtype), the histologic annotations (left) are shown with the cell deconvolution-based spot annotations (each spot is annotated as the dominant cell types from deconvolution results).

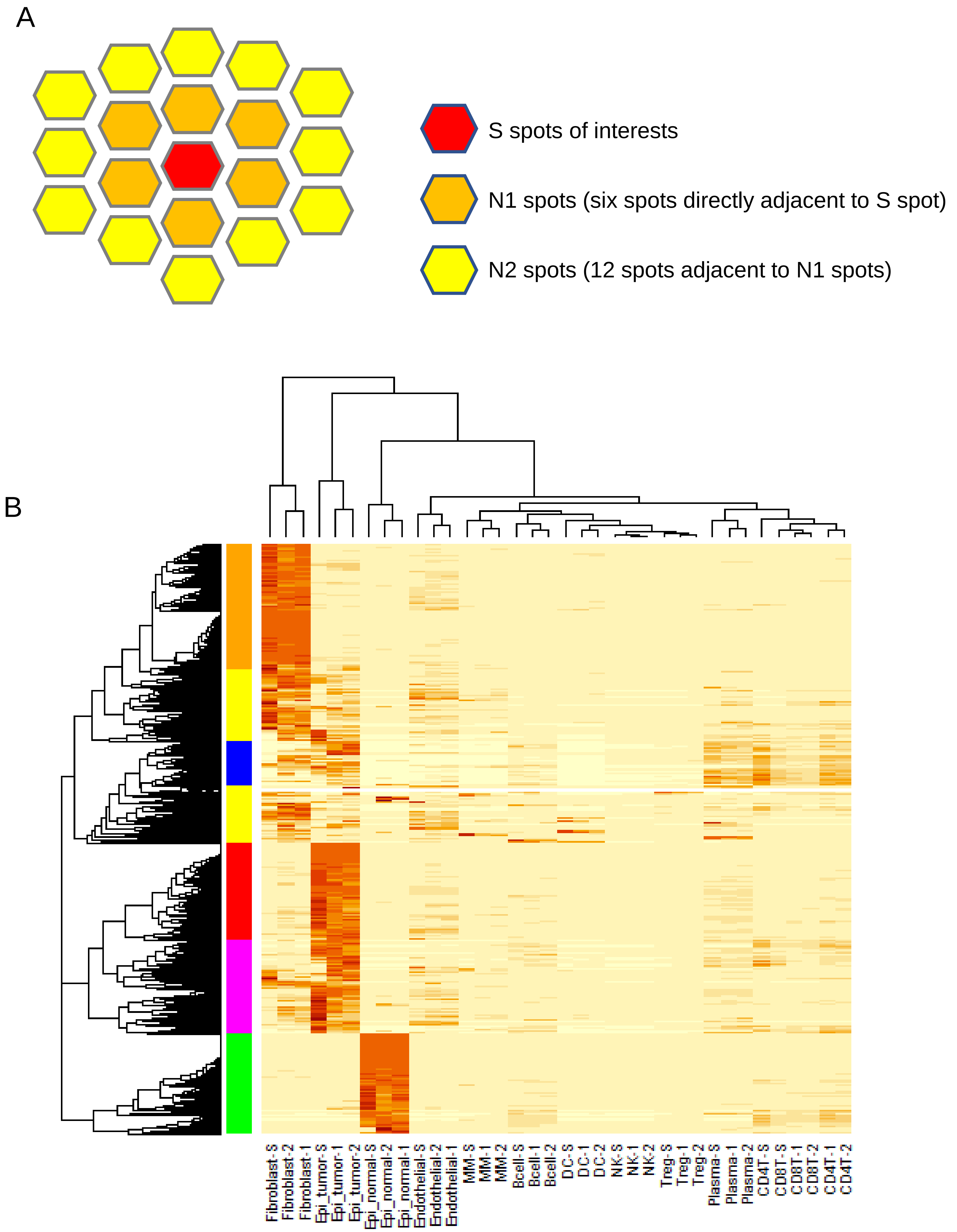


**Supplementary Fig. 2. Neighboring spot annotations and cellular abundance across all the spots observed.** (A) The spatial relationship of S/N1/N2 spots are shown in respective colors. (B) The abundance of 12 cell types for S (-S), N1 (-1) and N2 (-2) spots corresponding to main Figure 2A.

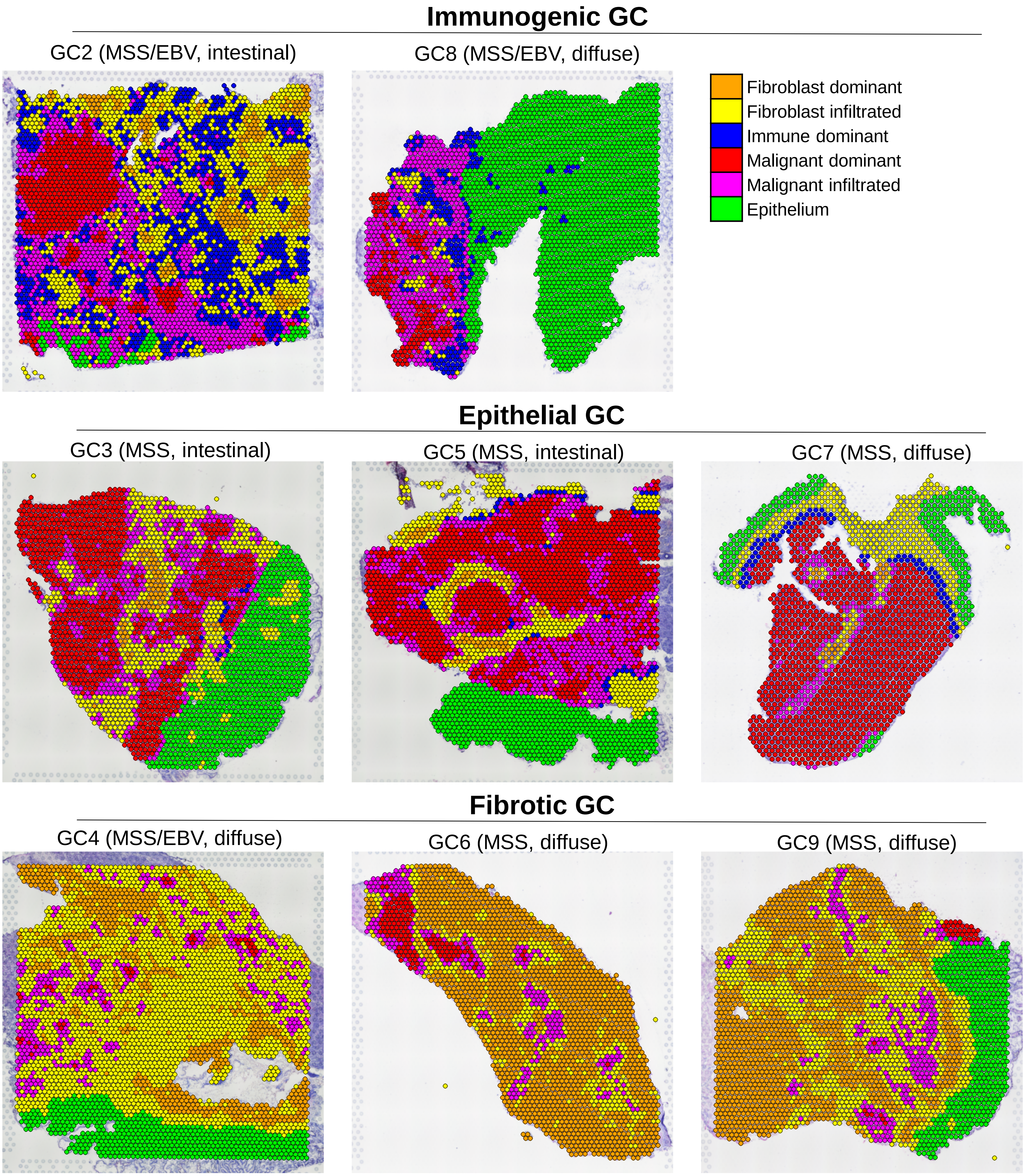


**Supplementary Fig. 3. CCZ presentation of GCs.** For the remaining cases (GC2-GC9), the spatial presentation of six CCZ are demonstrated illustrating the spatial architecture and relationship of CCZs.

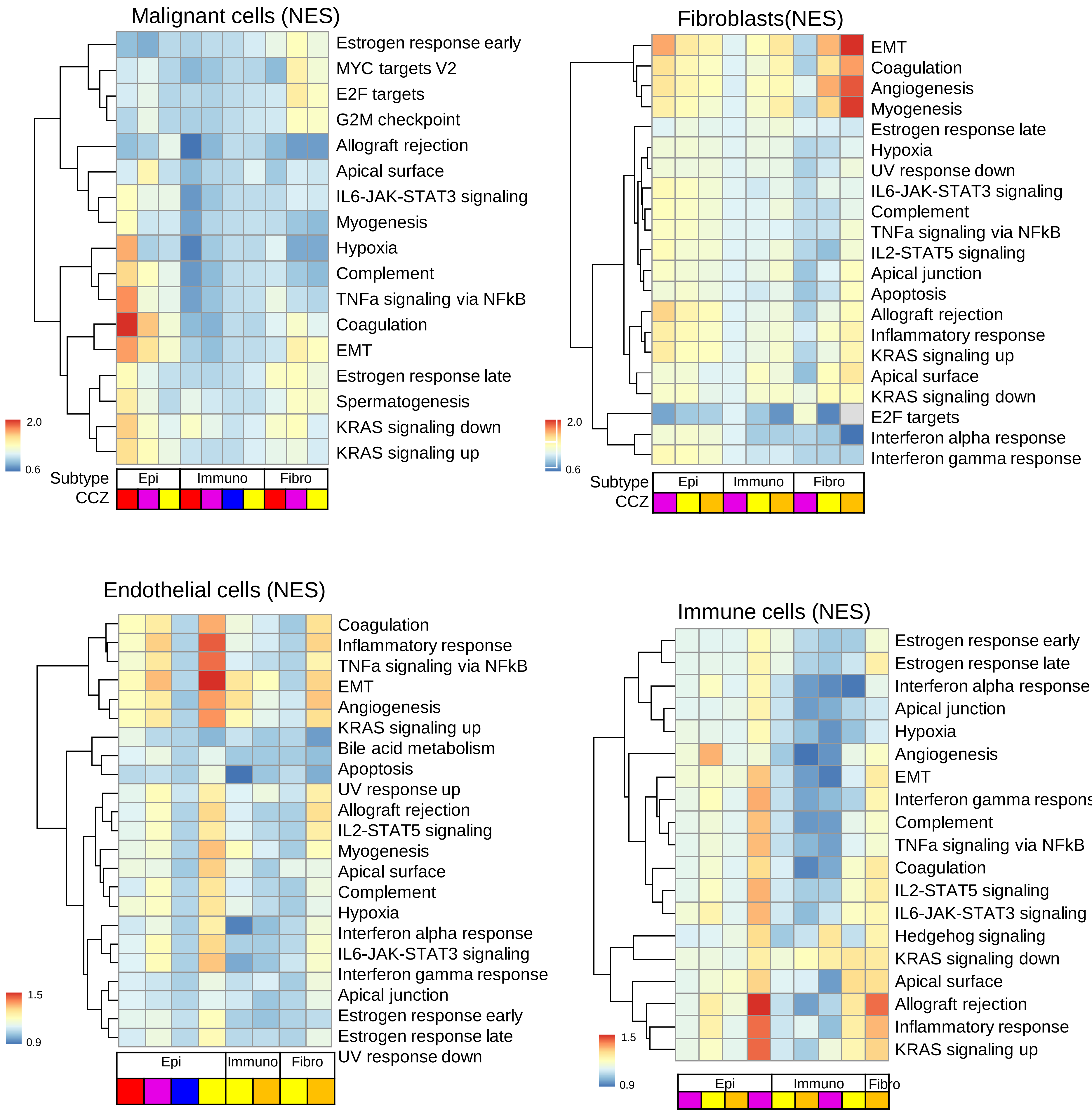


**Supplementary Fig. 4. Functional enrichment analyses of CCZ-specific cell type expression with respect to GC subtypes.** The functional enrichment analysis were done for CCZ-/cell type-specific expression that were inferred across GC subtypes are shown. GC subtypes and CCZs are colored below the heatmaps.

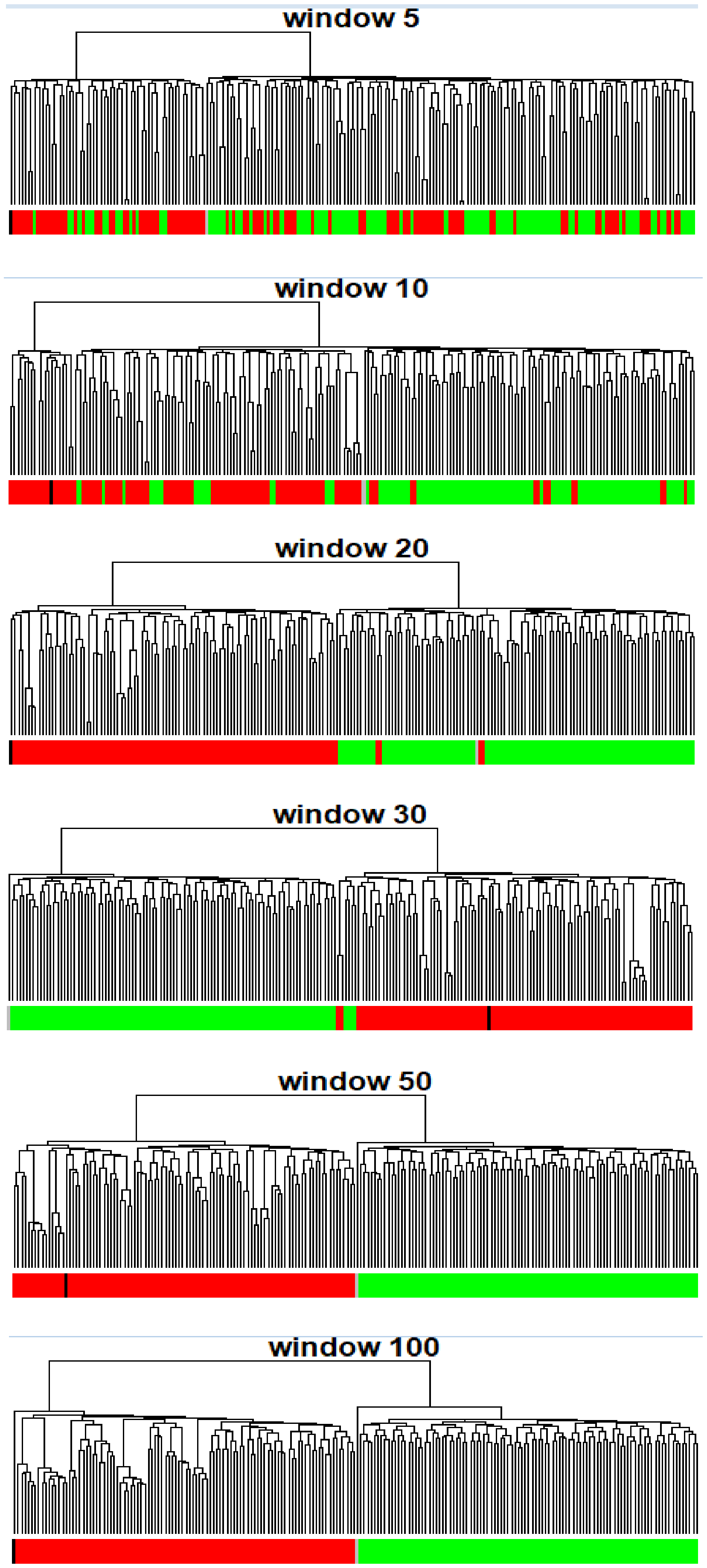


**Supplementary Fig. 5. Lineage concordance with respect to the number of windows or spot number.** Spots including 2 cell types were simulated and tested whether the expression of two cells deconvoluted across various number of spots (5 to 100) were tested.

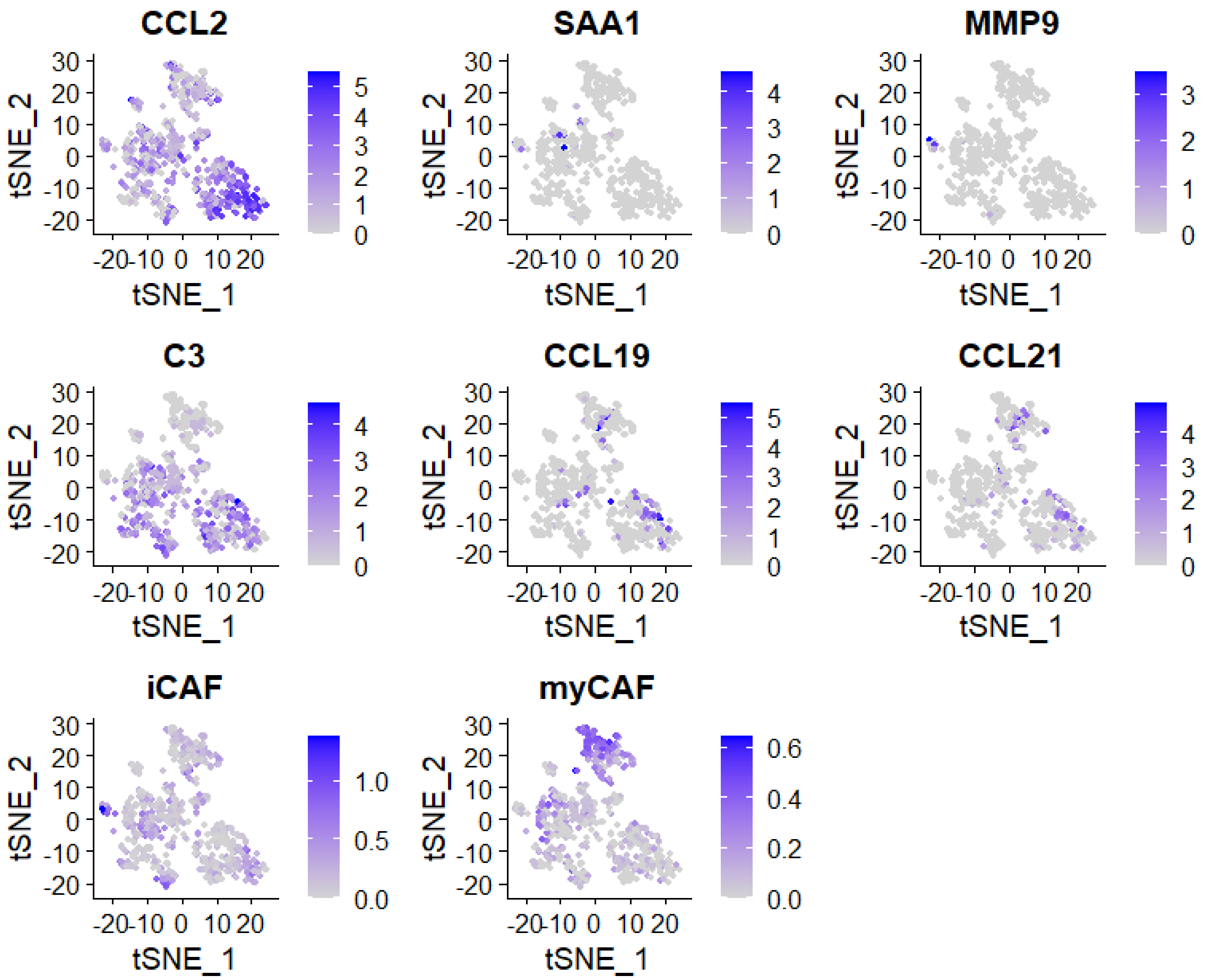


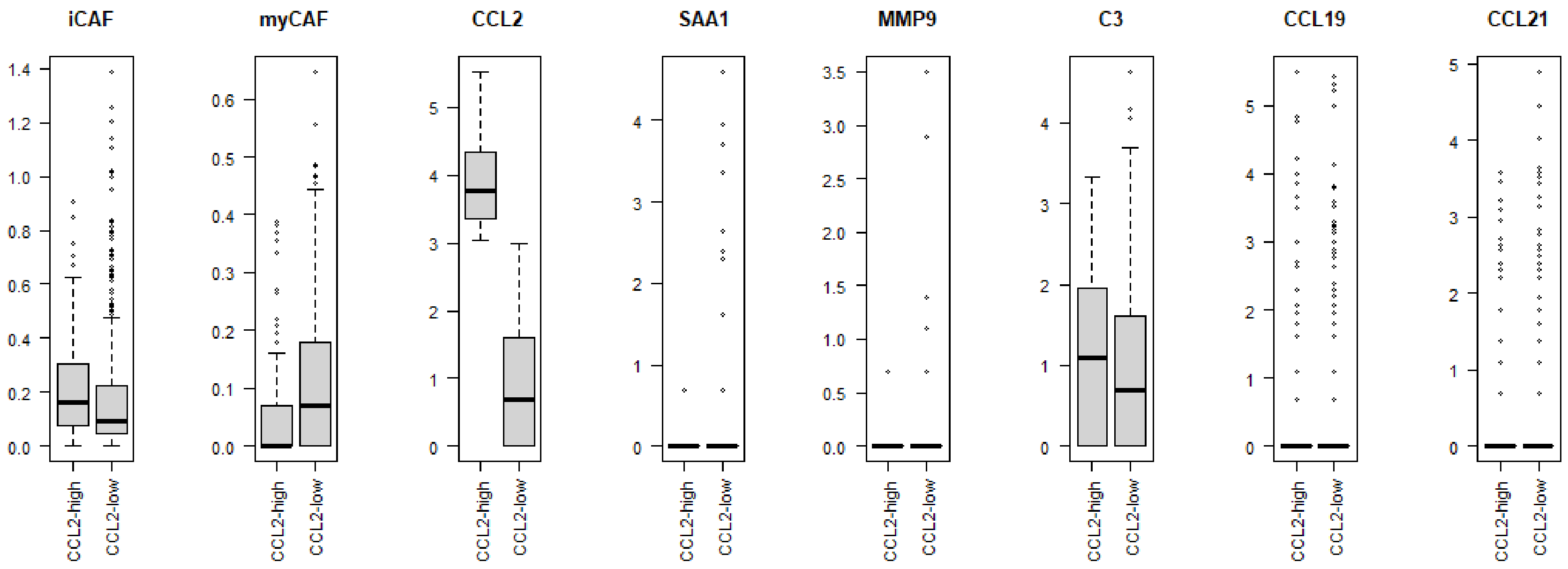


**Supplementary Fig. 6. CLL2+ fibroblasts and CAF markers.** Fibroblasts obtained from single cell RNA-seq data are shown for various ligand markers along with the scores of iCAF and myCAF in tSNE plots. Also shown for the differential level of the markers between *CCL2+* vs. *CCL2*-fibroblasts.

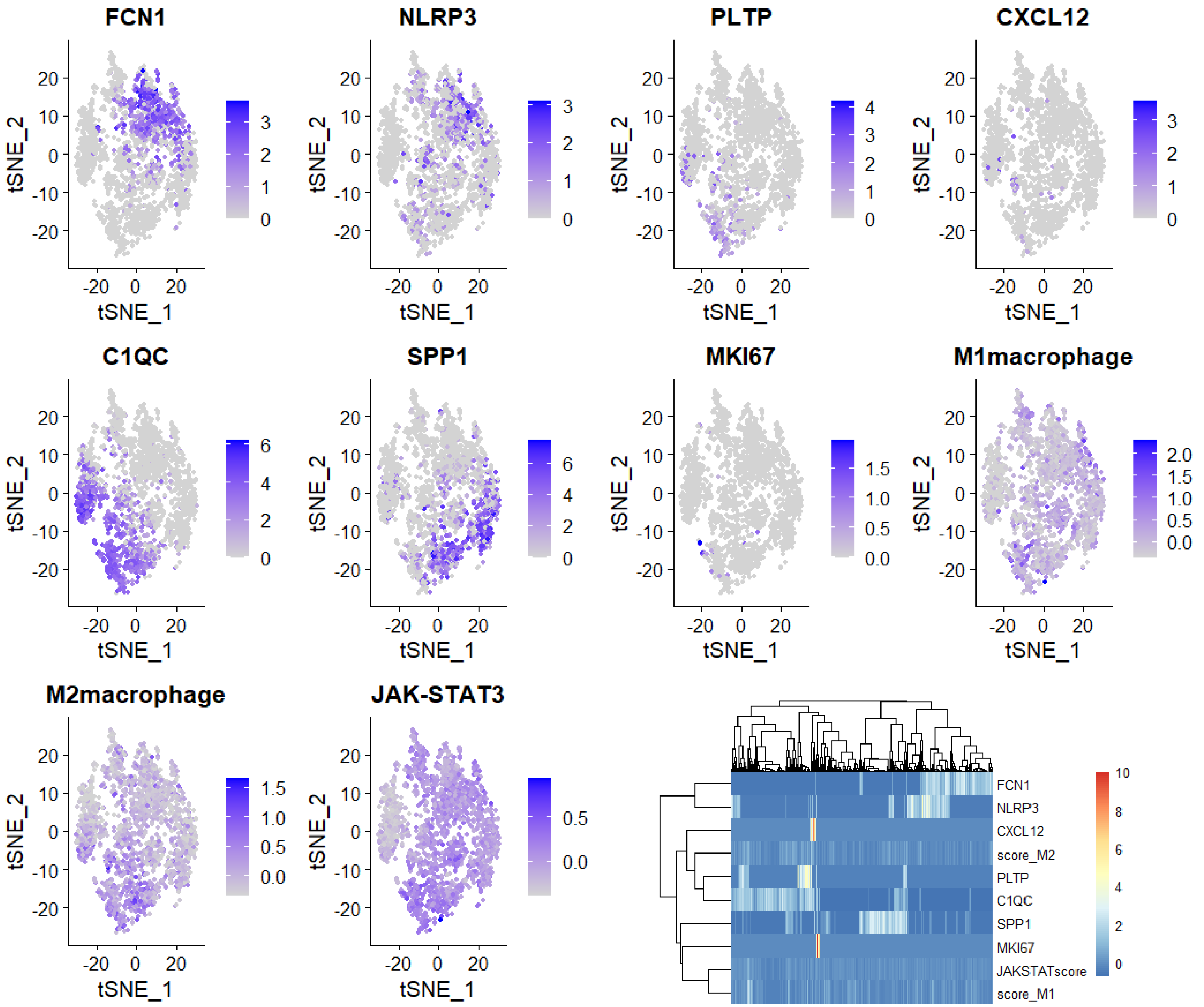


**Supplementary Fig. 7. Markers of macrophages.** Macrophagess obtained from single cell RNA-seq data are shown for various ligand macrophage markers along with the scores of M1/M2 macrophages and JAK-STAT3 signaling. Also shown a heatmap the relationship of the expression of the selected markers and scores.

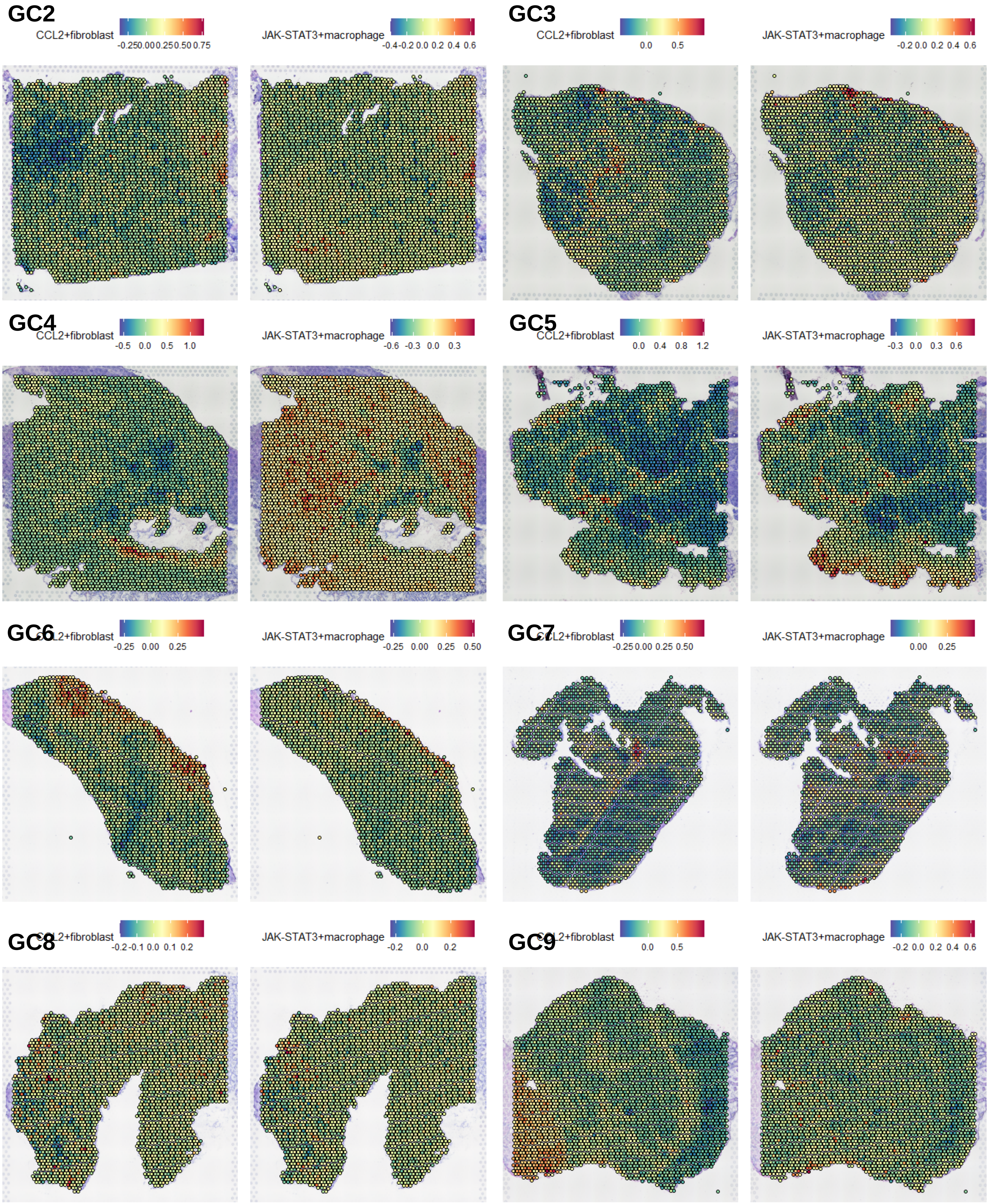


**Supplementary Fig. 8. Co-localization of CF (CCL2+fibroblast) and JM (JAK-STAT3+macrophages) abundance in spatial data**. For 8 cases (GC2 – GC9), the signature score-based abundance of CF and JM cells are shown in spatial data.

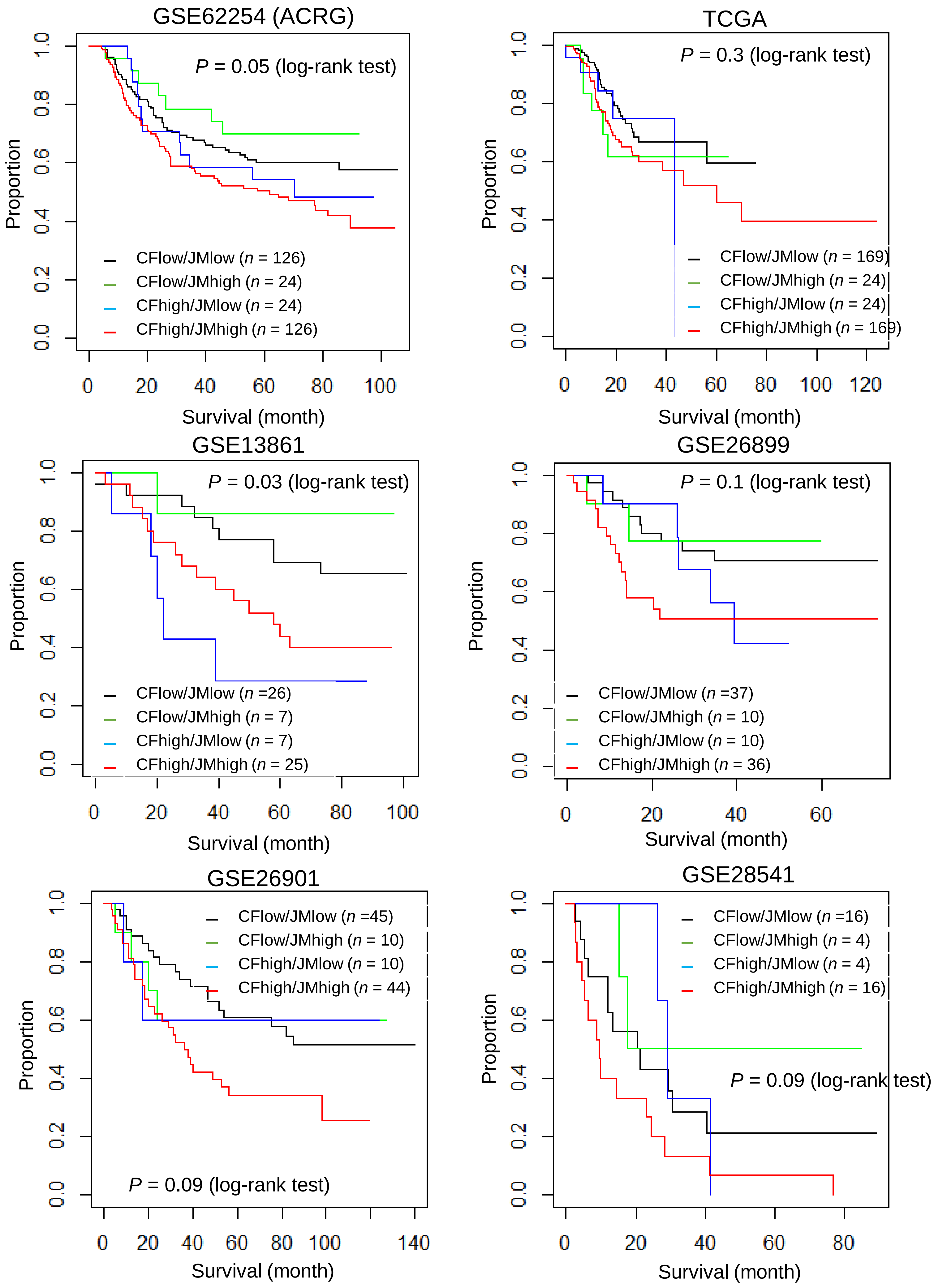


**Supplementary Fig. 9. Clinical outcome of CF (*CCL2*+fibroblast) and JM (JAK-STAT3+macrophages) abundance**. For six public database of GC bulk transcriptome with clinical outcome. Four categories of patients annotated by the abundance of CF and JM are compared for overall survival using KM-survival curves and log-rank tests.

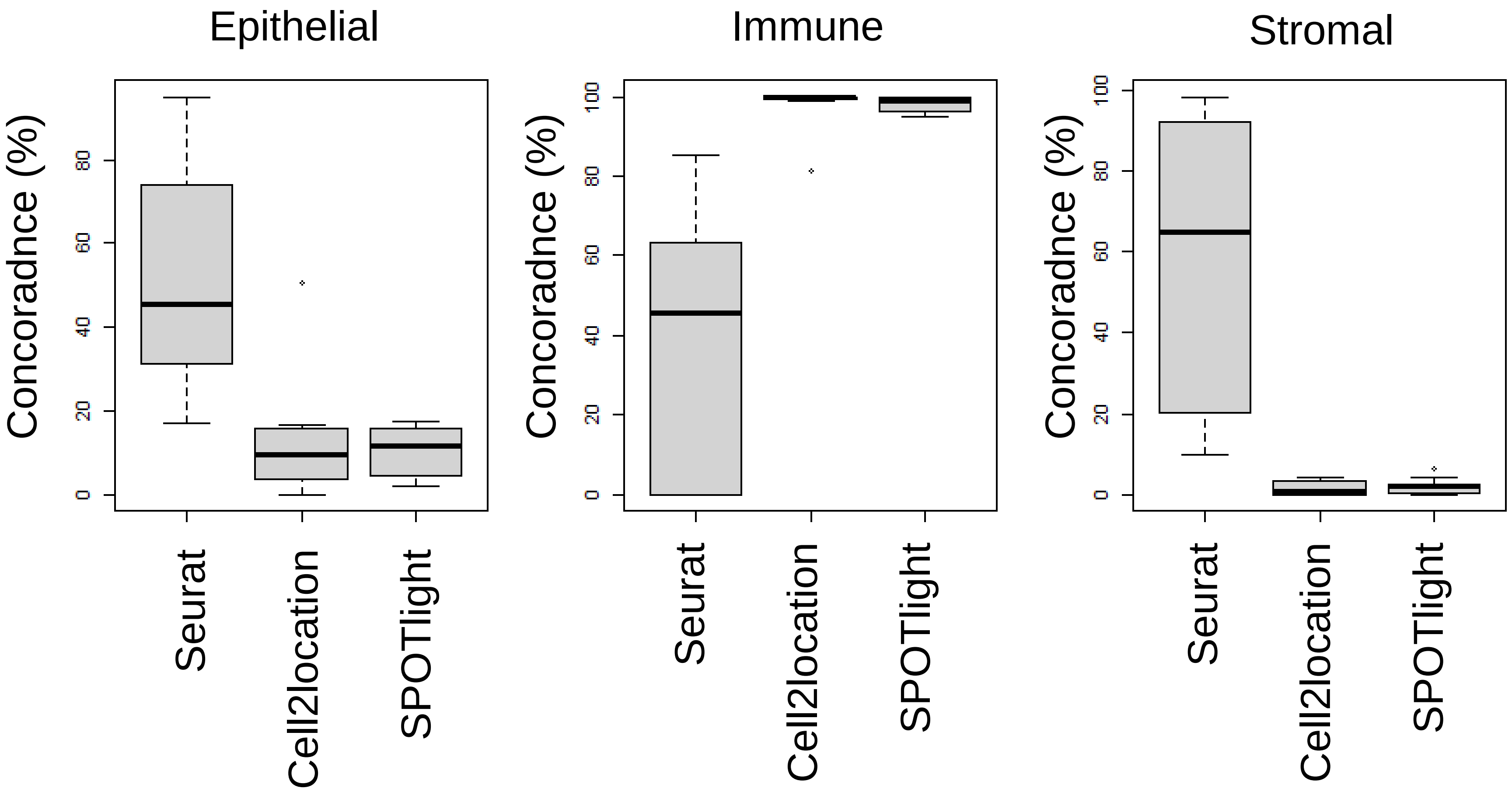


**Supplementary Fig. 10. Concordance of three deconvolution algorithms.** Three deconvolution algorithms of Seurat, cell2location and SPOTlight were compared for their concordance with histological annotations. Histological annotations were made by pathologist into three regional categories of ‘epithelial’, ‘immune’ and ‘stromal’. The comparisons were made for the histological annotations with corresponding cellular abundances deconvoluted by each of algorithms for 11 cell types.
